# Can Chemical Exchange Saturation Transfer (CEST MRI) be used as biomarker of disease progression in Prion Disease?

**DOI:** 10.1101/2021.12.16.473014

**Authors:** Eleni Demetriou, Mohamed Tachrount, Matthew Ellis, Jackeline Linehan, Sebastian Brandner, John Collinge, Simon Mead, Karin Shmueli, Mark Farrow, Xavier Golay

## Abstract

Human prion diseases are fatal neurodegenerative disorders that may have prolonged asymptomatic incubation periods. However, the underlying mechanism by which prions cause brain damage remains unclear. In turn, characterization of early pathological aspects would be of benefit for the diagnosis and potential treatment of these progressive neurodegenerative disorders. We investigated chemical exchange saturation transfer (CEST) MRI based on its exquisite sensitivity to cytosol protein content as a surrogate for prion disease pathology. Three groups of prion-infected mice at different stages of the disease underwent conventional magnetic resonance imaging and CEST MRI at 9.4T. For each mouse, chemical exchange contrasts were measured by applying five RF powers at various frequency offsets using magnetization transfer asymmetries. Relayed Nuclear Overhauser effects (NOE*) and amide proton transfer (APT*) were also assessed. For comparison, CEST MRI measurements were also made in healthy control mice brains. Here we show that alterations in CEST signal were detected before structural modifications or any clinical signs of prion disease. The detected CEST signal displayed different patterns at different stages of the disease indicating its potential for use as a longitudinal marker of disease progression. Highly significant correlations were found between CEST metrics and histopathological findings. A decline in NOE signal was positively correlated with abnormal prion protein deposition (R^2^ = 0.91) in the thalami of prion infected mice. Moreover, the NOE signal was negatively correlated with astrogliosis (R^2^ = 0.71) in the thalamus. No significant correlations were detected between NOE signals and spongiosis. MTR asymmetry at 3.5 ppm was also correlated with astrogliosis (R^2^ = 0.59), and prion protein deposition (R^2^ = 0.63) in thalamus. No significant changes were detected in APT* between prion-infected and control mice at all stages of the disease. Finally, MTR asymmetry between 2.8 and 3.2 ppm was correlated with prion protein deposition (R^2^ = 0.47) in the thalamus of prion -infected mice. To conclude, CEST MRI has potential utility as a biomarker of neurodegenerative processes in prion disease.

## Introduction

Human prion diseases are neurodegenerative disorders that progressively lead to lethal decline of cognitive and motor functions. The key event in the pathogenesis of prion disease is conversion of normal synaptic prion protein into the pathogenic state due to conformational changes occurring in the native protein structure. Human forms of prion disease include sporadic and familial Creutzfeldt-Jakob disease (CJD) affecting one to two people per million in the population worldwide annually, whilst for genetic forms such as Gerstmann-Sträussler-Scheinker (GSS) syndrome and Fatal Familial Insomnia (FFI) (1), the incidence is ten times less. In addition, there are several forms of transmitted prion diseases such as iatrogenic CJD, kuru, and variant CJD (1). A common feature of all these illnesses is their clinically silent incubation periods in which prion propagation rapidly reaches fixed levels of infectivity followed by the clinical onset of the disease (1). The lack of treatments for prion diseases makes the search for a disease-modifying intervention through clinical trials a research priority. There is a need to complement clinical outcome markers with quantifiable non-invasive biomarkers, in particular due to the absence of markers that might be able to report on disease pathology prior to or at the very beginning of symptoms onset.

To date, diagnosis of CJD can only be confirmed neuropathologically at autopsy or via brain biopsy. However, imaging methods are widely available and refined imaging sequences for early diagnosis of CJD would be highly desirable to complement existing biochemical tests such as determination of 14-3-3 protein in the CSF (2). Previous studies showed that MRI provides useful diagnostic information by showing microstructural features of deep and cortical grey matter structures that are found to be affected in Prion disease patients. Among different imaging sequences, Diffusion-Weighted Imaging (DWI) is considered the most sensitive diagnostic test (3). Early signs of microstructural changes in DWI scans were assigned to spongiform degeneration (4), misfolded prion protein accumulation (5) and neuronal loss (6). Another standard imaging sequence that is being applied in Prion disease patients is *T*_2_-weighted (*T*_2_w) imaging. High signal intensity has been observed in caudate, putamen and cortex (3) which was found to correlate with gliosis.

Chemical Exchange Saturation Transfer (CEST) or imaging of Nuclear Overhauser Effect (NOE) mediated by exchange-relayed signals or intermolecular pathways has been proposed as a new imaging mechanism to monitor protein folding *in vitro* by MRI (7). Changes in protein conformation are expected to alter the accessibility to water, which might affect slow chemical exchange processes between amide protons and water, and the associated altered peptide chain dynamics might also affect dipole-dipole intramolecular NOEs. In particular, reduced saturation transfer levels have been detected upon aggregation of amyloid beta and huntingtin proteins (8),(9) *in vitro* but there are, to our knowledge, no reports of chemical exchange imaging findings in prion diseases.

Therefore, the objective of our study was to determine the utility of CEST MRI to reveal specific pathologies in prion disease and establish that the underlying mechanisms of CEST make it a sensitive/specific biomarker for this disease. We compared CEST signals from different exchangeable proton groups found in proteins or larger macromolecules in the brains of prion-infected and control mice. To improve our detection sensitivity and selectivity we varied the irradiation amplitude to capture both slow and fast exchange processes. Once this was established, we performed mixed-effects regression analysis to assess whether our measurements showed significant alterations with progression of the disease and investigated the precision at which we could detect significant changes between the diseased and healthy mice. Finally, we correlated our imaging results with histopathological markers to report on the origin of the CEST signal in prion disease.

## Methods

### Protein study

Recombinant mouse prion protein was prepared and purified as described previously (10). Briefly, protein samples were converted from the native structure to the misfolded β-sheet conformation by reducing the protein disulfide bond and refolding at acidic pH (10). This was performed by denaturation of prion protein (PrP) in 6 M GdnHCl in the presence of 100 mM DTT to a final concentration of no more than 1 mg/ml, and subsequent refolding by dialysis against 10 mM sodium acetate, 2 mM DTT, pH 4. β-PrP^23–231^ was assembled into fibrils by treating 0.27 mg/ml of β-PrP^23–231^ in 10 mM sodium acetate, pH 4.0, with 1/9 volume of a 5 M stock of GdnHCl or NaCl (in the same buffer) to give final protein and denaturant concentrations of 75 μM.

All samples were imaged on a 9.4T Agilent scanner (Agilent Technologies, Santa Clara, CA, USA) at 20°C and 37°C using a transmit/receive RF coil with 33 mm inner diameter (Rapid Biomedical GmbH, Rimpar, Germany). CEST measurements were performed on a single slice using single-shot spin-echo (SE) echo planar imaging (EPI), (TR = 65.3 ms, TE = 4.07 ms, FOV = 20×20 mm^2^, slice thickness = 5 mm, matrix size= 64×64) with a saturation train prior to the readout consisting of 300 Gaussian pulses and at an irradiation amplitude of 1.5 μT (pulse length = 20 ms, FA = 982°, 95% duty cycle).

### Animal studies

#### Control and prion-infected mice

Work with mice was performed under licence granted by the UK Home Office and conformed to institutional guidelines. Two groups of 7-week-old female FVB mice were intracerebrally inoculated with 30μl of 1% brain homogenate of the Rocky Mountain laboratory (RML) prions train (n=19) or with brain homogenate from uninfected mice (Mock inoculum) as controls (n=11). The prion-infected group was separated into three groups of mice scanned at different stages of prion disease: 80 days post injection (dpi) – asymptomatic-stage (n=6); 130 dpi – early-stage (n=6); and 160 dpi – late-stage (n=7). Control mice were separated into two groups: 80 dpi (n=5) and 160 dpi (n=6). All mice were anaesthetized (1.5-1.8% isoflurane in 1.5 l/min oxygen with balance in air) and scanned on a 9.4 T Agilent system (Agilent Technologies, Santa Clara, CA, USA) using a 33-mm-diameter transmit/receive coil (Rapid Biomedical GmbH, Rimpar, Germany).

### CEST protocol

All images were collected at a single slice (thickness=2mm), centred on the thalamus. CEST MRI measurements were acquired using a turbo flash readout (matrix: 64×64, TR=2.11ms, TE=1.07ms, FOV=20×20mm2) preceded by a train of Gaussian pre-saturation radiofrequency (RF) pulses. Those pulses were applied at 71 frequency offsets to collect an evenly sampled Z spectrum between −5.0 and 5.0ppm. To obtain optimal saturation efficiency of slow and fast exchanging species, five different radiofrequency field amplitudes (B_1_) were applied (11),(12): 0.6μT (n=80, pulse length=50ms, flip angle=360°, 90% duty cycle), 1.2 μT (n=80, pulse length=50ms, flip angle=900°, 90% duty cycle), 2.0 μT (n=60, pulse length=50ms, flip angle=1200°, 90% duty cycle), 3.6 μT (n=40, pulse length=50ms, flip angle=2100°, 90% duty cycle) and 10 μT (n=30, pulse length=50ms, flip angle=6000°, 99% duty cycle). In addition, to improve image SNR and to reduce the physiological noise during MRI acquisition, each Z-spectrum was collected three times and then averaged for each mouse.

### CEST Data analysis

NOE* is mainly observed in the upfield frequencies, i.e., negative frequency offsets from water, and originates from dipole-dipole interactions between water and protons from the side groups of large mobile molecules such as aliphatic components of peptides, proteins and lipids. In this study, NOE* values were calculated between −3.7 and −3.3 ppm by finding the difference between the fitted Z-spectrum and a linear interpolation (13). Amide proton transfer (APT) is a dominant saturation effect in brain Z-spectra downfield (in the positive frequency range) from the water peak. Its major contributors are mobile proteins and peptides via exchangeable amide protons. APT* values were calculated between 3.3 and 3.7 ppm in a similar fashion to NOE*.

MTR asymmetries were calculated as the difference in signal on either side of the water peak for the frequency offsets listed in Supplementary Table 1. The effect of the B0 fluctuation was corrected by estimating the frequency offset corresponding to the minimum of a fitted spline (13). For each mouse, Regions-of-interest (ROIs) delimiting the cortex and the thalamus were drawn on the anatomical image and then applied on the corresponding NOE* and MTR asymmetry maps.

### Histological protocol

After the MRI, the mice were terminated by CO_2_ overdose and thirty mouse brains (five groups) were submitted for histological evaluation. Brains were fixed in 10% buffered formalin, and coronal slices of the brain were processed through graded alcohols (70, 90%, 100%) and xylene followed by paraffin wax (LEICA ASP 300; Leica Milton Keynes, UK), and embedded in wax blocks. The middle piece was embedded face down (see Supplementary Figure 1 A). Paraffin sections were cut using a Leica RM2235 microtome to a nominal thickness of 4μm. Five serial sections from each mouse brain were mounted on Superfrost™ microscope slides, air-dried and melted at 60°C for 30 minutes before staining.

A haematoxylin and eosin (H&E) stain was carried out on the first serial section from all the processed blocks using a Gemini AS autostainer (ThermoScientific) to identify the region of interest as shown in Supplementary Figure 1 (between A and B). The slides identified as containing the region of interest were then stained further with antibodies against abnormal synaptic prion protein ISCM-35 (ICSM35), glial fibrillary acidic protein (GFAP) for astrocytes, and ionized calcium-binding adaptor protein-1 (Iba1) for microglia according to manufacturers’ guidelines.

### Histopathological analysis

All histological slides were digitised using a Leica SCN400 (LEICA Milton Keynes, UK) producing 8-bit, brightfield, whole slide images (WSI) at a resolution of 0.25 μm per pixel. These WSI were then analysed using Definiens Developer version 2.3 (Definiens AG, Munich, Germany). Following identification of the tissue region, the thalamus, hippocampus and cortex were manually delineated. The RGB image was then converted to huesaturation-density (HSD) representation (14), giving a measure of brown/red and blue stain intensity at each pixel.

For ICSM35, GFAP and Iba1 stained sections the chromogen distribution was quantified: All brown pixels were identified where the brown intensity was >0.1 arbitrary units (au) higher than blue intensity. Brown objects were then further segmented to Low Intensity <= 0.4au < Medium Intensity <= 0.8au < High Intensity. The area and relative coverage of these intensity classes were then exported for the three regions of interest.

Vacuolisation (spongiosis) was quantified on haematoxylin and eosin (H&E) stained sections: Nuclei were identified as regions with blue intensity greater than the average intensity of blue in the tissue region. Vacuoles were identified as regions with red intensity lower than the average intensity of red in the non-nuclear tissue; these were then “grown” where red and blue intensity was below a dynamically defined threshold. Any potential vacuole that shared a border with a nucleus was excluded as a shrinkage artefact; any that shared a border with glass was excluded as background; and any with an area less than 5μm^2^ or greater than 5000μm^2^ or with an elongated shape were excluded as non-vacuole spaces. The area, number, relative coverage and density of vacuoles were then exported for the three regions of interest.

### Statistical analysis

NOE* and APT* values were assessed for significance between prion-infected and control mice using t-tests where significant changes are indicated by p<0.05.

Differences in MTR asymmetry measures among prion-infected mice scanned at all stages of the disease were assessed using mixed-effects regression models in STATA (15). A mixed-effects analysis allowed the examination of multiple variables simultaneously for predicting an outcome measure of interest. In our case the variables considered were the MTR asymmetries calculated at various irradiation amplitudes and frequency offsets in all mice brains for both thalamus and cortex. In the case of NOE* and APT* we performed t-test analysis because we only consider a single range of frequencies and one irradiation amplitude. The question we addressed was whether our measurements showed substantial alterations with the progression of the disease. The relationship between MTR asymmetries, NOE*, APT* and histopathological scores was assessed across the whole group of prion infected mice with Spearman’s rank correlation. In all cases, significant changes are indicated by p<0.05. Mean and standard error of the mean of all measures are reported as mean ± standard error.

## Results

In the first part of the study, indication of prion protein misfolding was assessed in samples containing native and misfolded prion protein (PrP) by collecting CEST spectra as described above. Additionally, T2-weighted images in the brains of prion-infected mice were visually inspected and quantitative NOE*/APT* maps were used to assess pathological features originating from altered prion protein conformation and accumulation of misfolded PrP.

### In vitro studies of native and misfolded prion protein

Figure 1 (a-b) shows the Z-spectra and the MTR asymmetry lineshapes obtained from native (blue) and misfolded (red) prion protein at a concentration of 75 μM and at a pH of 7.2. Z-spectra of the misfolded protein display reduced saturation transfer levels at 3.5 ppm and 1.8 ppm corresponding to amide and amine exchanging resonances as well as reduced exchange-relayed NOE between −2.0 and −5.0 ppm. These changes could be attributed to the increased content of β-sheet structures in the misfolded prion protein. Native prion protein was also scanned at pH 6.4 (red) and 7.2 (blue) and at 20 °C and 37 °C (see Supplementary Figure 2) to assess changes in the measured CEST contrast due to pH and temperature.

**Figure 1.**
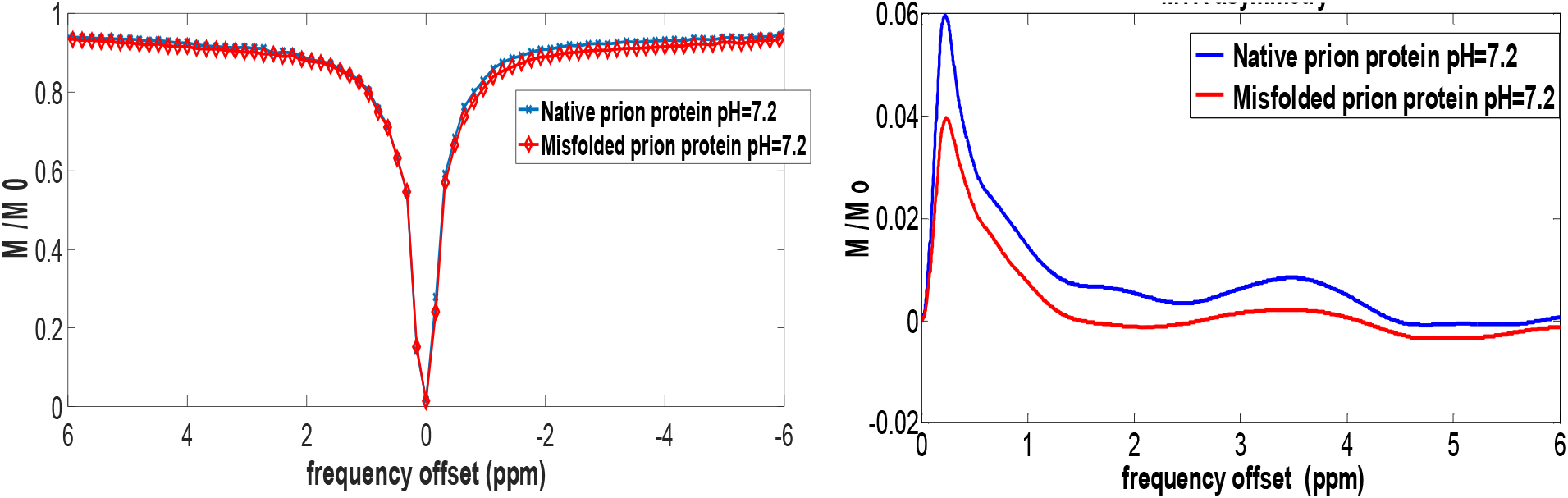
Z-spectra and MTR asymmetry lineshapes of native and soluble β-sheet-rich oligomers of prion protein (75 μM) at a pH of 7.2 and at room temperature.

Overall, an increase in CEST contrast was observed at higher prion protein concentration and higher pH (see Supplementary Figure 2 (e), (f)).

### In vivo studies

#### Structural changes in Prion disease

Figure 2 shows conventional T_2_ weighted images obtained from the brains of prion-infected mice at 80 dpi, 130 dpi and 160 dpi and control mice at the first and last timepoints. Increased grey matter signal intensity was observed in 80 dpi prion-infected mice relative to controls in cortical and hippocampal regions. These hyperintensities in the T_2_w images at 80 dpi in the prion-infected mice could be related to the reaction of the brain to the injection of prion-containing brain homogenate and might not be directly related to the progression of the disease in this model.

**Figure 2.**
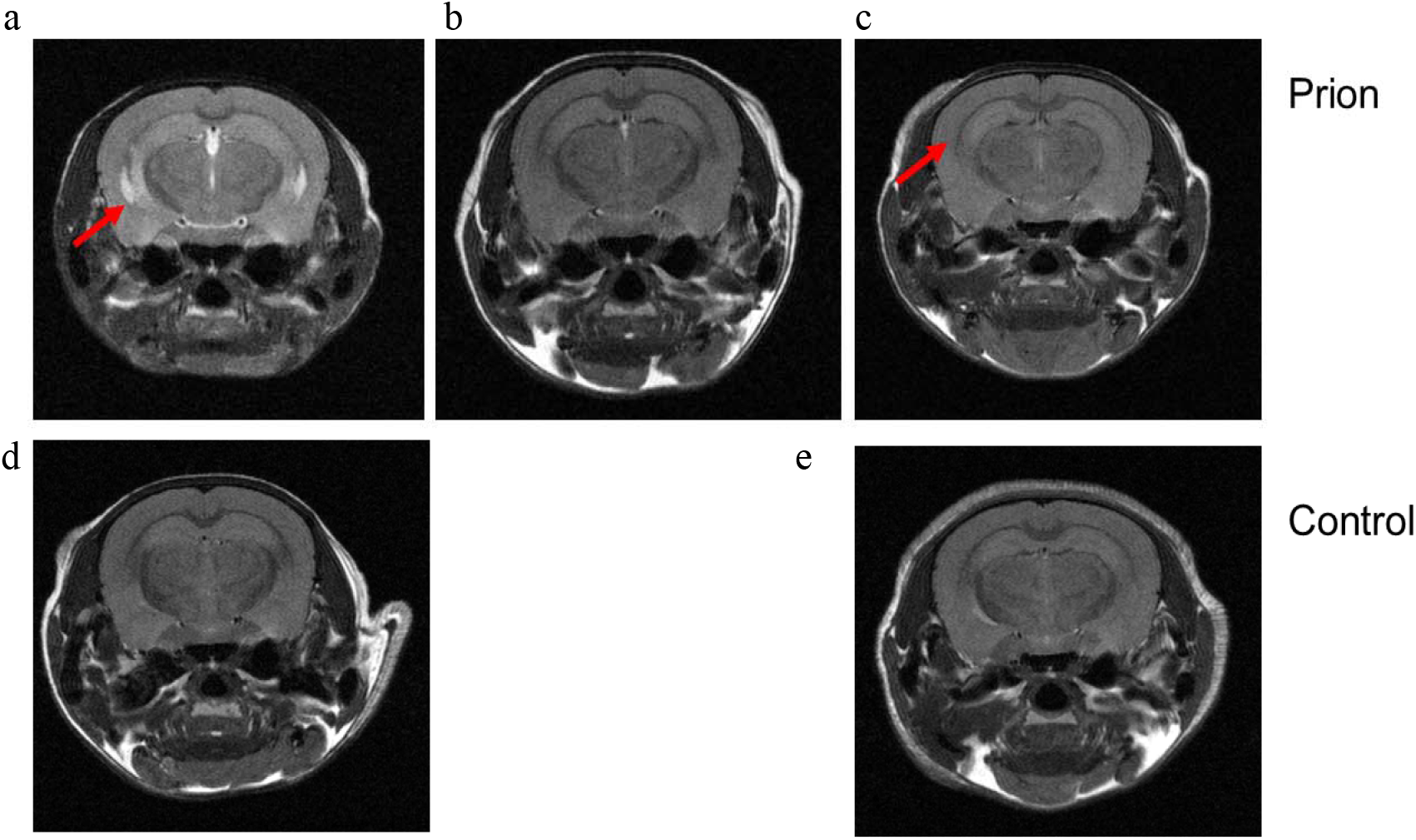
Structural changes in (a) asymptomatic, (b) early-stage and (c) late-stage prion-infected mice at the level of the thalamus, (d-e) Control mice brains at matched time points. Red arrows indicate hyper-intensities in T2w images of prion infected mice.

### NOE*, APT* analysis

NOE* and APT* concentrated on the frequency of chemical exchange e.g. −3.5 ppm or 3.5 ppm while the contrast from surrounding frequencies (e.g. −3.7 to −3.3 ppm) was used as a baseline for magnetization-transfer-free NOE* and APT* contrast. In the case of NOE*, a significant decrease (p<0.05) was observed in the thalamus and cortex of prion-infected mice relative to healthy controls throughout the disease course. No significant difference (p=0.12) was detected between the two control groups scanned at 80 dpi and 160 dpi, and they were merged into one group of mice when compared with prion-infected mice. NOE* was significantly reduced in the thalamus and cortex of prion-infected mice at 130dpi (p=0.00008 for thalamus, p=0.001 for cortex) and 160dpi (p=0.00005 for thalamus, p=0.003 for cortex) when compared with the control group (see Figures 3 and 4). Furthermore, NOE* was significantly reduced (p=0.001) in the thalamus of asymptomatic mice (80dpi) but there was no significant change in the cortex (p=0.14). Significant reduction of NOE* in the cortex of prion-infected mice was observed only at 130dpi and 160dpi. This is consistent with observations that abnormal prions show a global pattern of deposition at later stages of the disease, while at 80dpi pathological changes are found mostly in the thalamus and brainstem, not in the cortex (16). No significant changes in APT* were detected between prion-infected and control mice at any disease stage possibly indicating the lack of pH changes in prion mice as well as no alteration in the proteasome content (Supplementary figure 3).

**Figure 3.**
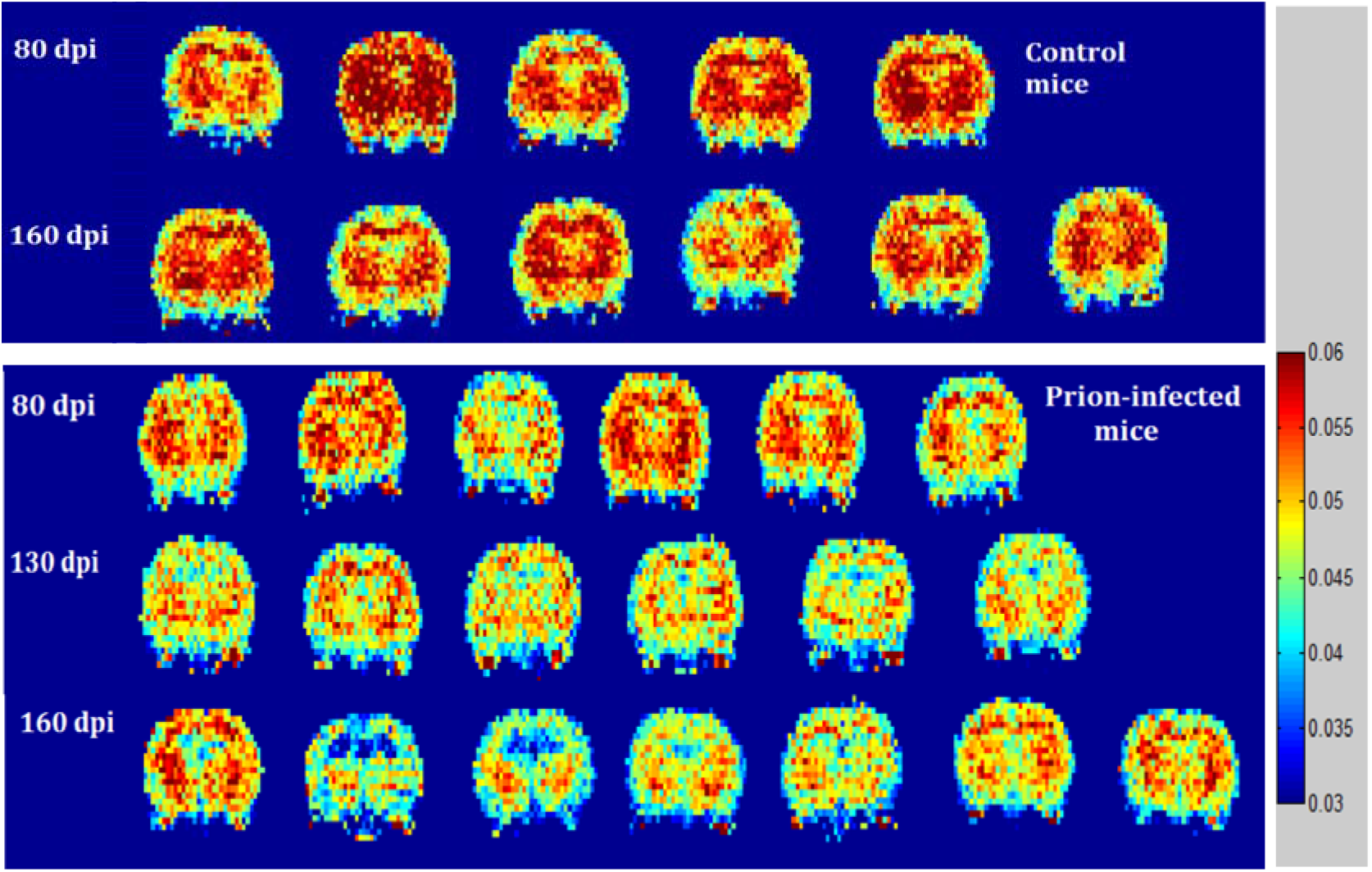
NOE* maps for control mice (top) and prion-infected mice (bottom) at different stages of Prion disease. NOE* values were found to be significantly reduced in the brains of prion-infected mice when compared with the controls.

**Figure 4.**
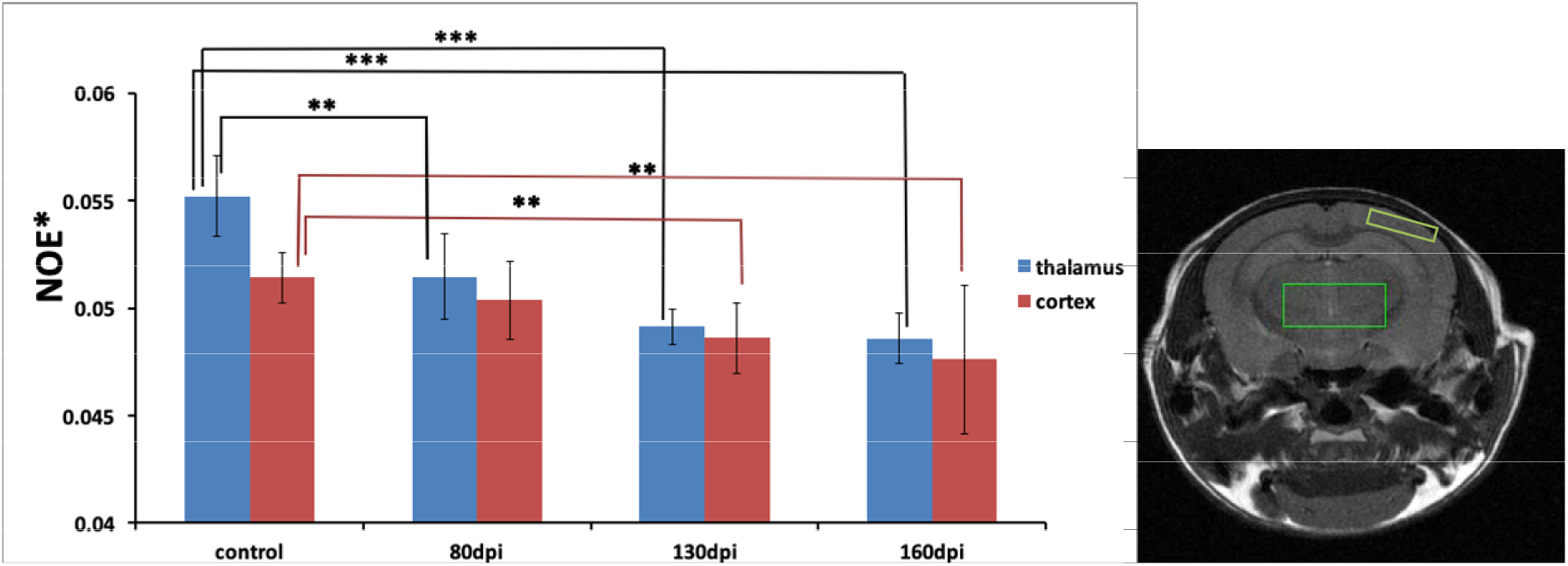
NOE* between −3.7 and −3.3 ppm in thalamus and cortex of prion-infected mice at different stages of prion disease and control mice (a). Two-tailed t-tests were used to evaluate changes in NOE* values measured in prion-infected and control mice (** p <0.01, *** p < 0.001). The thalamic (green) and cortical (yellow) ROIs are shown overlaid on an anatomical image (b).

### MTR analysis

In the second part of the study, CEST signals from all mice were calculated with MTR asymmetry analysis at various irradiation amplitudes and frequency offsets. To investigate disease progression, a two-tailed t-test analysis was performed (mice were separated into five groups). Regional alterations in CEST signal were assessed in the cortex and the thalamus of prion-infected and control mice.

#### 80 dpi prion-infected mice vs 80 dpi controls

Significant differences were found between the asymptomatic prion-infected mice vs control mice for all frequency offsets at 10 μT in the cortex. Moreover, a significant increase was detected at 2.8-3.2 ppm and 1.0-1.5ppm in the thalamus. As expected, significant changes in CEST signal of asymptomatic mice were less severe than the signal changes observed in the late stage of prion disease. Among the 80 dpi prion-infected group we were able to detect two subgroups of mice with more (n=4) or mild (n=2) prion pathology according to the MTR asymmetry measurements. Supplementary Table 3 displays the significant changes in MTR asymmetry of asymptomatic prion-infected and control mice in the thalamus and cortex which shows that the MTR asymmetry of prion-infected mice is smaller compared to the control group of mice. Representative Z-spectra obtained from the thalamus and cortex of an asymptomatic and a control mouse are shown in Supplementary Figures 12 and 13.

#### 160 dpi prion-infected mice vs 160 dpi controls

MTR asymmetries of prion-infected mice were less negative compared to the MTR asymmetries of the control group in the thalamus at 0.6 μT and 1.2 μT. In the cortex, the MTR asymmetries decreased for the prion-infected mice compared to the control group. This trend was visible for high irradiation amplitudes greater than 2.0 μT. Supplementary figures 4–8 summarise the calculated MTR asymmetries from all mice in both the cortex and thalamus and for all irradiation amplitudes and frequency offsets used. Supplementary Table 2 displays the significant changes in the MTR asymmetry observed in prion-infected and control mice in the thalamus and cortex. Representative Z-spectra obtained from the thalamus and cortex of a late stage prion-infected mouse at five irradiation amplitudes are shown in Supplementary Figures 9–11. Z-spectra from a control mouse at 160 dpi are also displayed for comparison (SI Figures 9–11).

### Progression of the pathology in prion-infected mice

#### Prion-infected mice: 80 dpi versus 130 dpi

Reduced MTR asymmetries were detected in thalamus and cortex of 80 dpi prion-infected mice when compared to 130 dpi prion-infected mice (see Supplementary Table S4 in SI).

#### Prion-infected mice: 130 dpi versus 160 dpi

No significant changes were detected when comparing the MTR asymmetries of 130 dpi prion-infected mice with those in 160-dpi prion-infected mice.

#### Prion-infected mice: 80 dpi versus 160 dpi

Reduced MTR asymmetries were detected in 80 dpi prion-infected mice when compared to 160 dpi prion-infected mice (see Supplementary Table S5 in SI) at a few frequency offsets in the thalamus. In addition, higher MTR asymmetry was detected at 2.8-3.2 ppm in the cortex. Supplementary Table S5 displays the comparison of 80 dpi vs 160 dpi prion-infected mice. Supplementary Figures S14 and S15 display the averaged Z-spectra from 80 dpi, 130 dpi and 160 dpi prion-infected mice at five different irradiation amplitudes.

MTR data was imported into STATA and significant differences in MTR asymmetries between control and diseased mice were assessed using a mixed-effects model analysis. In particular, MTR asymmetries from all mice in both cortex and thalamus and for all frequency offsets were evaluated as dependent parameters while the irradiation amplitude was treated as an independent variable.

The results obtained using mixed-effects model analysis further confirm significant differences in MTR asymmetry between prion and control mice at low irradiation amplitudes: 0.6 μT, 1.2 μT as well as at high powers such as 3.6 μT and 10 μT in both the thalamus and the cortex of prion infected mice (p< 0.05). A summary of the results is displayed in Supplementary table 7.

### Histological results

Thirty brains were analysed for spongiform alterations by haematoxylin and eosin (H&E) staining together with immunohistochemical analyses for deposition of abnormal PrP, astrocytic gliosis, and microglia activation. Figure 5 shows pathological markers in prion-infected brains at 80 and 160 dpi. Figure 6 shows the levels of abnormal prion protein, microglial activity and astrogliosis in prion-infected and control mice.

**Figure 5.**
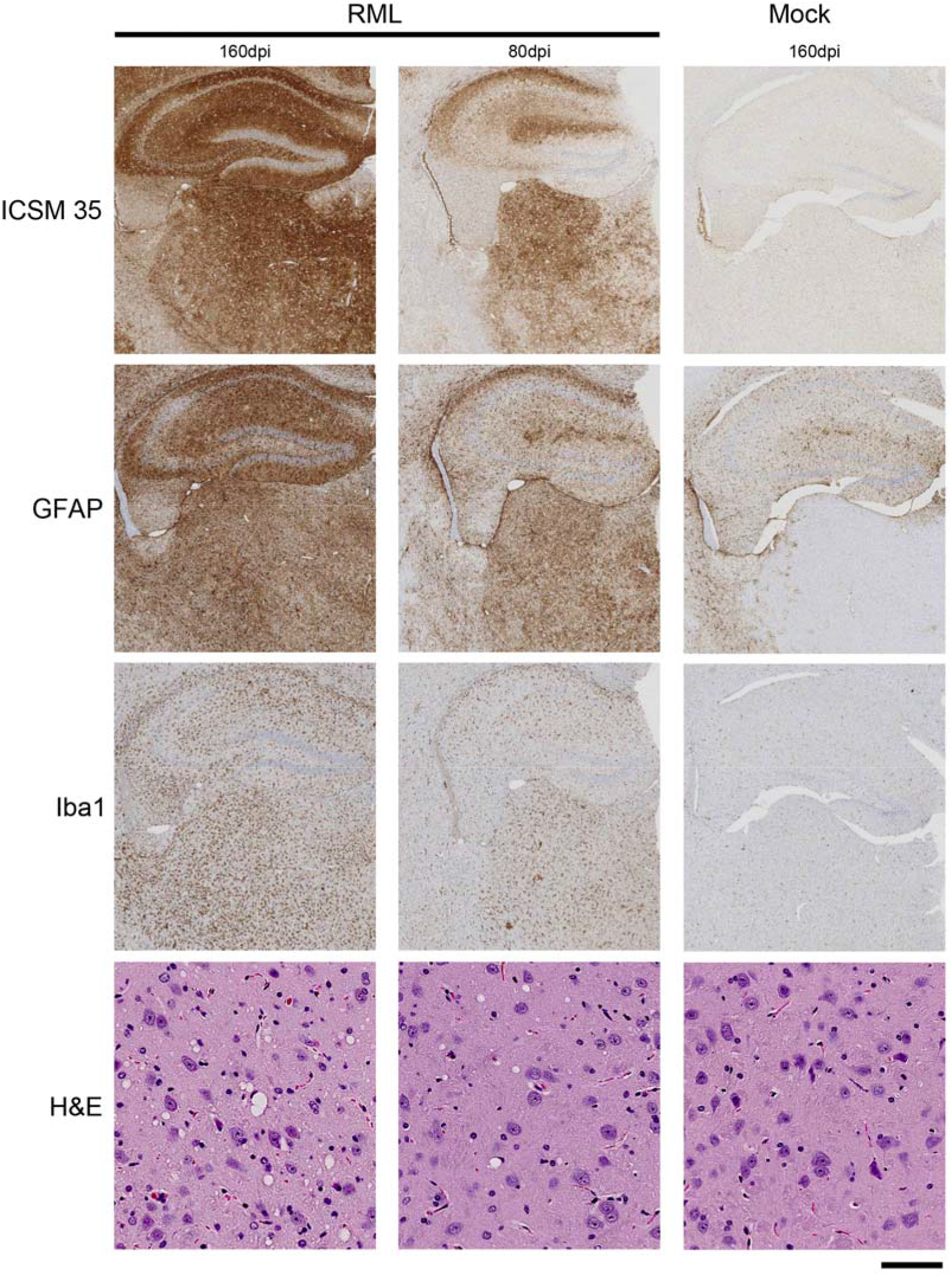
Demonstration of prion protein deposition (ICSM35), astrogliosis (GFAP) and microglia activity (Iba1) in the hippocampus and thalamus and spongiosis (H&E) in the striatum of RML infected mouse brains at 80 dpi and 160 dpi and controls at 160 dpi. Scale bar=100 **μm** and 1mm.

**Figure 6.**
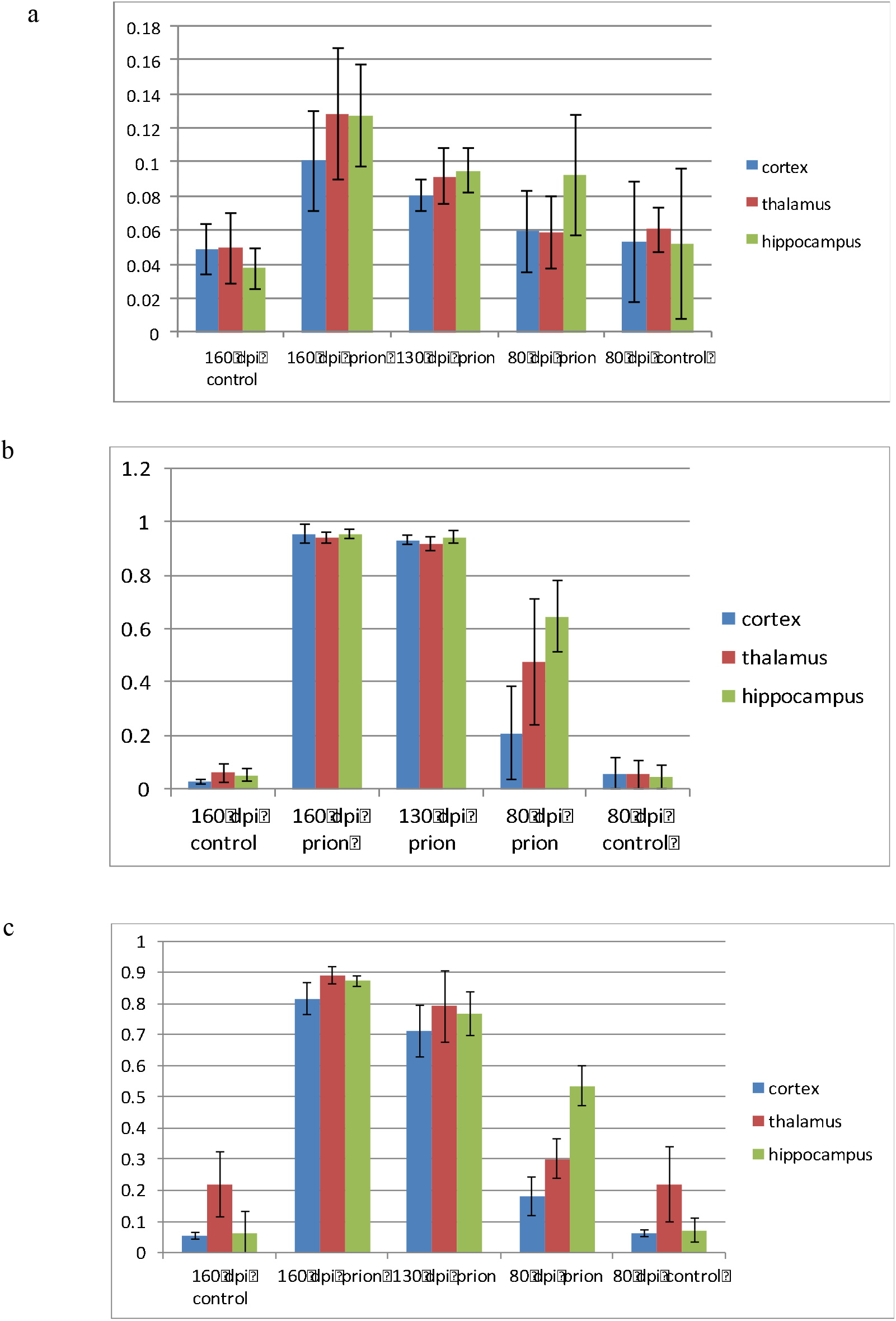
Levels of microglia activation (a), abnormal prion protein (b) and astrogliosis (c) in healthy mice and prion-infected mice in cortex, thalamus and hippocampus.

The experimental group of 80 dpi prion-infected mice displayed mild general pathology, considerably less than the experimental group of 130 dpi. Pathological changes include widespread moderate deposition of synaptic prion protein (ICSM35) in thalamus and hippocampus, visible overall activation of reactive astrocytes but much less than in group 130 dpi and mild-to moderate microglial activation in hippocampus. Focal spongiosis was detected in hippocampus, mild spongiosis in thalamus and none in cortex.

Overall, the experimental group of 130 dpi prion-infected mice showed strong involvement of all regions but less than the experimental group of 160 dpi prion-infected mice. The pathological changes include widespread, moderate to severe, deposition of synaptic prion protein (ICSM35), strong widespread astrogliosis with accentuation in the hippocampus and thalamus, strong activated microglial reaction in thalamus and hippocampus and widespread spongiosis throughout the brain (less severe compared to 160 dpi).

The experimental group of 160 dpi prion-infected mice displayed severe prion pathology consistent with the end-stage of prion disease. Pathological changes include widespread strong deposition of synaptic prion protein (ICSM35), astrogliosis, spongiform degeneration in grey matter structures and activated microglia cells in thalamus and, to a lesser extent, in hippocampus. The cortex was less affected with a mild accentuation of prion protein, gliosis, spongiosis, and microglia pathology in the deep cortical layers towards the corpus callosum. Prion protein deposition covers all brain regions with very little difference between individual regions. In addition, there is a slight predominance of hippocampal and thalamic gliosis compared to cortical regions. Spongiosis was found to be moderate to severe in the cortex and severe in the hippocampal and thalamic regions.

### Correlation of histological findings with CEST results

The relationship between CEST measurements and histological findings was assessed using Pearson correlation in all diseased mice. In thalamus, MTR asymmetry at 3.5 ppm correlated with astrogliosis (R^2^=0.59), prion protein deposition (R^2^=0.63) and microglia activity (R^2^=0.37). MTR asymmetry between 2.8-3.2 ppm was also correlated with prion protein deposition (R^2^=0.47) and astrogliosis (R^2^=0.32) in the thalamus of prion infected mice. These correlations are all associated with increased MTR asymmetries in prion mice versus healthy controls. In the cortex, no significant correlations were detected between prion-infected and control mice. No correlations were found for prion protein or astrogliosis, the absence of which yielded lower MTR asymmetries in prion mice versus controls. In thalamus, NOE* was correlated positively with abnormal PrP (R^2^= 0.93) and negatively with astrogliosis (R^2^= 0.71). Finally, misfolded PrP and astrogliosis were positively correlated (R^2^= 0.93).

## Discussion

In this study, we used a mouse model of prion disease to investigate if CEST could be used as an early biomarker of protein metabolic dysfunction. Aggregation of prion protein in mice is characterised by deleterious changes in the central nervous system including neuronal degeneration, vacuolation and gliosis. Disease specific signs in mice are well characterised in terms of the development of pathological features throughout the disease course based on histological markers (16). Prion propagation occurs in two stages: an exponential phase (phase 1) which reaches steady levels of infectivity irrespective of the concentration of the normal prion protein, followed by a linear rise (phase 2) of disease-related prion protein (16). The clinical and neuropathological features of prion disease are detected only when phase 2 is well established. According to our imaging results, a few significant changes were detected in asymptomatic mice, followed by global changes in NOE* and MTR asymmetry analysis as the disease progressed in agreement with pathological markers seen in prion-infected mice.

The first question we aimed to answer was whether the CEST technique is sensitive enough to detect a difference in signal solely due to prion pathology in mice. Decreased levels of NOE* were detected in the thalamus of prion-infected mice at all stages of the disease, but only at the early and late stages in the cortex, which correlated with abnormal PrP deposition (R^2^= 0.93). While our *in vitro* results from native and misfolded prion proteins show a similar decrease in NOE signal, there is a problem of scale of the effect. Indeed, at −3.5 ppm, the difference in signal was found to be 0.00525 au in the solution of misfolded vs native protein compared with 0.00625 au between prion and control mice. Yet, from private communications, a realistic concentration of the misfolded PrP in prion-infected mouse brain is around 0.286 micromolar which is two orders of magnitude lower than the *in vitro* concentration we scanned. As such, it is unlikely that the aspect of PrP misfolding could play a direct role in the CEST signal we are detecting.

However, evidence suggests that toxic β-sheet isoforms inhibit the ubiquitin-proteasome system (17). This pathway is important for the rapid elimination of misfolded cell proteins and many other proteins critical in regulating gene expression and metabolism (17). In prion mice increased levels of ubiquitin conjugates have been identified, which correlate with decreased proteasome activity. It is possible that soluble micro-aggregates of misfolded proteins overwhelm the ubiquitin system causing a functional impairment. Histological analysis confirms the accumulation of the misfolded isoform in the brains of mice even at the asymptomatic stage in thalamus which then progressed in other critical brain regions. Similarly, NOE* follows a global reduction in the brain as the disease progresses which could be linked to a decreased proteasome activity because of the presence of the aggregated PrP rather than the direct detection of the misfolded protein in mice brains. NOE* signal was correlated with both astrogliosis and accumulation of misfolded PrP. There is evidence that astrocytes are the cell type in which the abnormal form of the prion protein is first replicated in the nervous system (17). It is likely that the changes in astrocytes induced by PrP^Sc^ are due to the “stress” induced in these cells by the peptide.

Considering MTR asymmetry analysis, at the late stage of the disease, the MTR asymmetries were found to be significantly larger in diseased relative to control mice at 0.6 μT in thalamus. Considering that the NOE peak is a very broad peak, covering resonance frequencies between −1.0 and −5.0 ppm, the less-negative asymmetries we measured in prion-infected mice can be interpreted either as a reduction in the NOE peak area or as an additional increase at positive-frequency offsets. Furthermore, the MTR asymmetries in the cortex of late-stage prion-infected mice were found to be significantly decreased, compared to the control mice. This trend was visible for B_1_ > 2.0 μT, in which magnetisation transfer contrast from macromolecules becomes more pronounced. Clinically, the magnetisation transfer ratio was found to be reduced in both the whole brain and the grey matter of symptomatic Prion-disease patients (18). In addition, MTR decline in formalin-fixed cerebral and cerebellar hemispheres of Prion-disease patients has been shown to be negatively correlated with spongiosis scores obtained from histology (18). Here, however, while MTR asymmetry at 3.5 ppm correlates mainly with astrogliosis and prion protein deposition in thalamus there were no positive correlations in the cortex of prion mice. Therefore, we hypothesise that the positive MTR asymmetries in prion disease might be associated with neuronal degeneration secondary to the aggregation and insolubility of the misfolded prion protein, or to other molecular pathologies as discussed above. It is worth noting that the amplitude of the frequency-selective pulses is tuned to probe specific exchangeable species based on their characteristic exchange rates. In our study, we varied B_1_ in a non-linear fashion to cover almost all the exchange processes happening *in vivo* as shown in our previous work in rats undergone stroke (19).

The progression of prion disease in mice was investigated by comparing CEST results at three different stages of the disease. Significant changes were detected, mainly in the cortex of asymptomatic prion-infected mice; the MTR asymmetry was found to be reduced when compared to control mice. At the end stage of the disease a global reduction in the MTR asymmetry was observed in the cortex and a global increase in the thalamus when compared to age-matched controls. Furthermore, when comparing the early-stage prion-infected mice with asymptomatic mice, a global significant increase in MTR asymmetry was observed in the thalamus of early-stage prion-infected mice, and only at a few frequency offsets in the cortex. Interestingly, the comparison between asymptomatic and late-stage prion-infected mice showed less significant changes in MTR asymmetries relative to the comparison between asymptomatic and early-stage prion-infected mice. A possible explanation for this could be that the developmental changes that take place during the period from 80 days to 160 days in the brains of all mice alter the baseline CEST signals in the same way as the progression of prion disease. Therefore, some of the significant changes between prion-infected mice are cancelled out.

In a recent study, CEST was used as a marker of neurodegeneration in a transgenic mouse model of Alzheimer’s disease. Elevated CEST signals were detected in the brains of diseased mice and were correlated with higher glial cell proliferation, compared to wild type mice (20), while focusing on myo-inositol, which resonates at 0.6 ppm. However, the data displayed a global increase in the CEST signal compared to control mice. In human clinical studies, elevated MTR asymmetries were also detected in Parkinson’s disease patients (21) and in MS patients (22). The origin of higher CEST signals in neurodegenerative diseases is still unclear and it is hypothesised to be associated with the accumulation of abnormal cytoplasmic proteins such as α-synuclein in Parkinson’s Disease. Finally, reduced glutamate weighted CEST signals at 3.0 ppm were reported in a mouse model of Huntington disease (23). Despite these efforts, it should be mentioned that, even at fields as high as 9.4T mapping of individual metabolites such as glutamate is challenging mainly because of the overlap with other coalesced resonances such as taurine and glutamine. Thus, interpretation becomes more difficult. Here, we reported changes across a larger range of frequencies using MTR analysis and, although a molecular interpretation might not be clear, the global changes in MTR which we reproducibly measured in mice show promise for translation to clinical practice.

### Limitations

One of the shortcomings of this study was that we did not include a group of control mice at 130 dpi and therefore it was difficult to assess whether there are any changes in CEST signal associated with developmental changes at this timepoint.

## Conclusion

The findings of this study suggest the use of CEST as a non-invasive imaging technique to probe neuropathological changes in prion-infected mice. The larger changes (detected at the end stage of the disease) are in line with behavioural and histological results. We hypothesise that the measured signal changes could be attributed to decreased proteasome activity because of the accumulation of the aggregated PrP rather than the direct detection of the misfolded prion protein in mice brains. Alterations in the CEST signal were detected before any structural changes or clinical signs of prion disease which is encouraging in terms of the ability to translate CEST techniques to aid in the early detection of prion disease and other neurodegenerative disorders.

## Acknowledgements

We would like to thank Dr Emmanuel Risse for preparation of prion protein solutions. We are grateful to MRC technical staff for animal handing during the experiment and Dr Laslzo Hosszu for insightful discussions. We would like to thank Prof Allan Hackshaw and Dr Michael Katsoulis for their input and feedback on mixed effects model analysis. We would like to thank Franscisco Torrealdea and Marilena Rega for insightful discussions and Andreia Silva for assistance during the experiment. This project was funded by UCL Grand Challenge (ED), and by the Department of Health’s NIHR Biomedical Research Centre’s funding scheme to UCL/UCLH (SB, XG, MT). It has also received funding from the European Union’s Horizon 2020 research and innovation programme under grant agreement No 667510 (ED, XG).

---

**Supplementary Figure 1:**
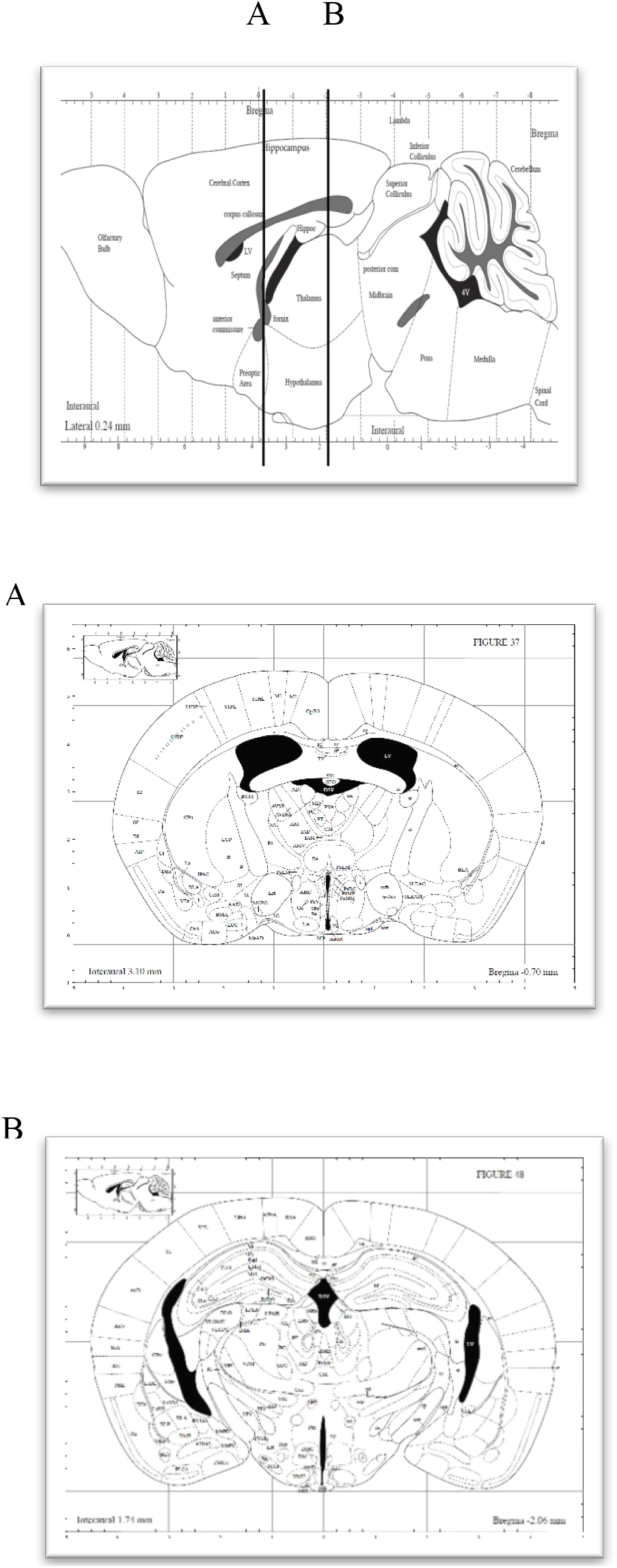
Sagittal diagram shows the coordinates of the obtained paraffin sections. All coronal sections were analysed from the anterior hippocampus (Bregma −0.70 mm, A) to the mid-posterior hippocampus (Bregma −2.06 mm, B).

**Supplementary Figure 2:**
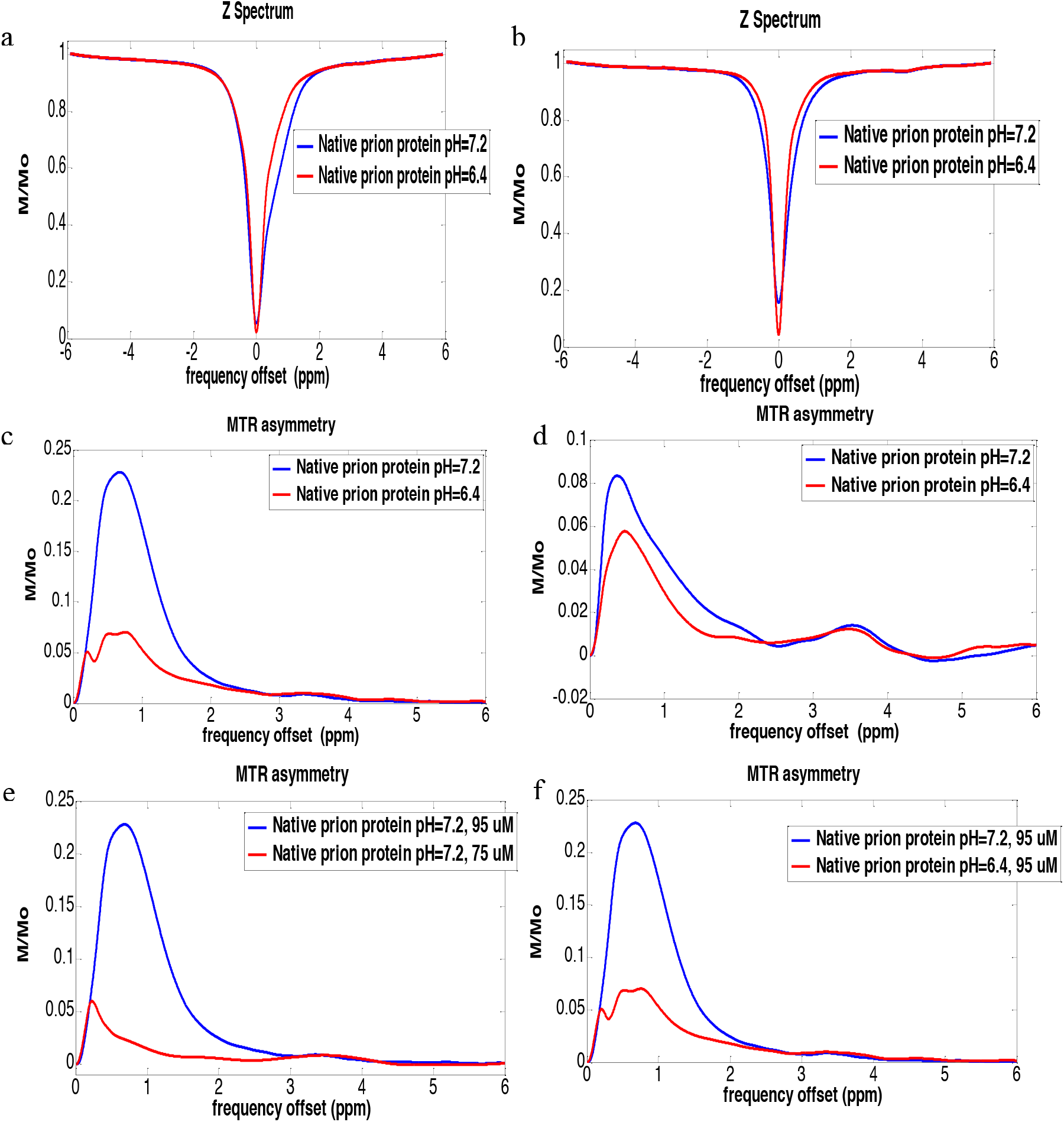
Z-spectra and MTR asymmetry lineshapes for native prion protein at pH 6.4 and 7.2 at 37 °C (a,c) and 20 °C (b,d).

The contrast at resonance frequencies between 0.2-2.0 ppm (assigned to hydroxyl and amine groups) was larger for pH 7.2 and for both temperatures when compared to pH 6.4. (see Supplementary Figure 2). In addition, amide proton exchange at 3.5 ppm is visible on the Z-spectra collected at 37 °C. Overall, an increase in CEST contrast is observed at higher prion protein concentration and higher pH (see Supplementary Figure 2 (e), (f)).

**Supplementary figure 3:**
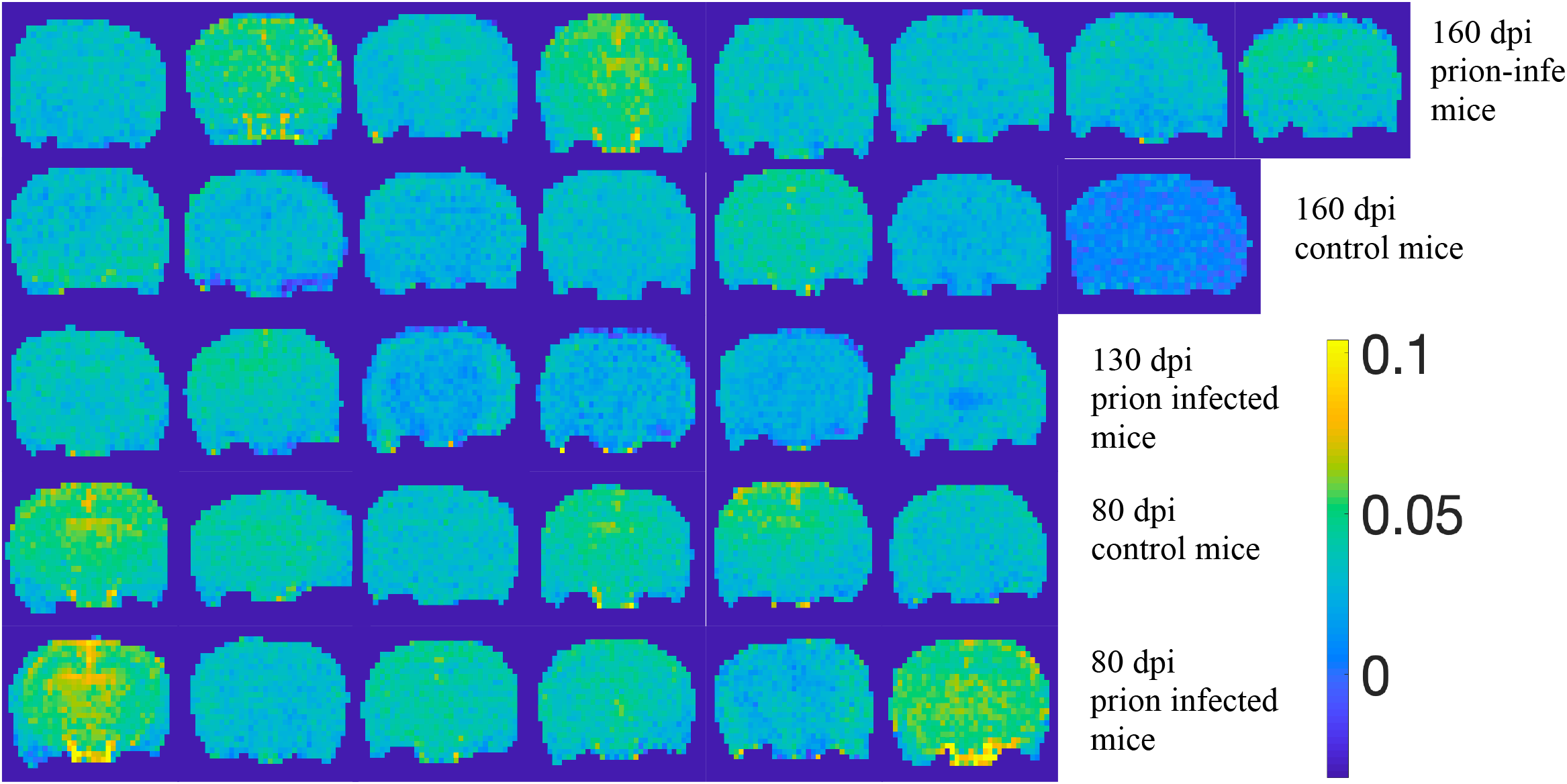
APT* maps in prion infected and control mice: (a) 160 dpi control mice, (b) 160 dpi prion infected mice (c) 130 dpi prion infected mice, (d) 80 dpi control mice, (e) 80 dpi prion mice

**Supplementary Table 1.**
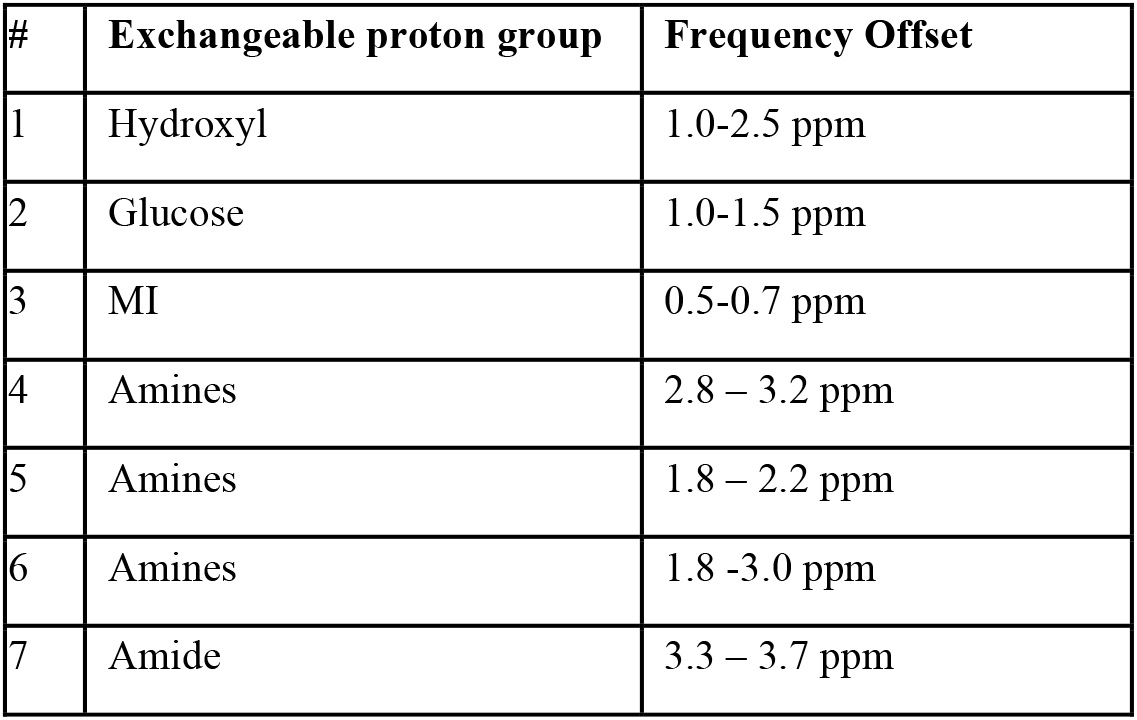
displays the proton resonances for various exchangeable proton groups as found in the literature. MTR asymmetry analysis has been performed to assess whether prion neuropathology affects the dynamics of molecular processes resonating at distinct frequency offsets.

**Supplementary figure 4:**
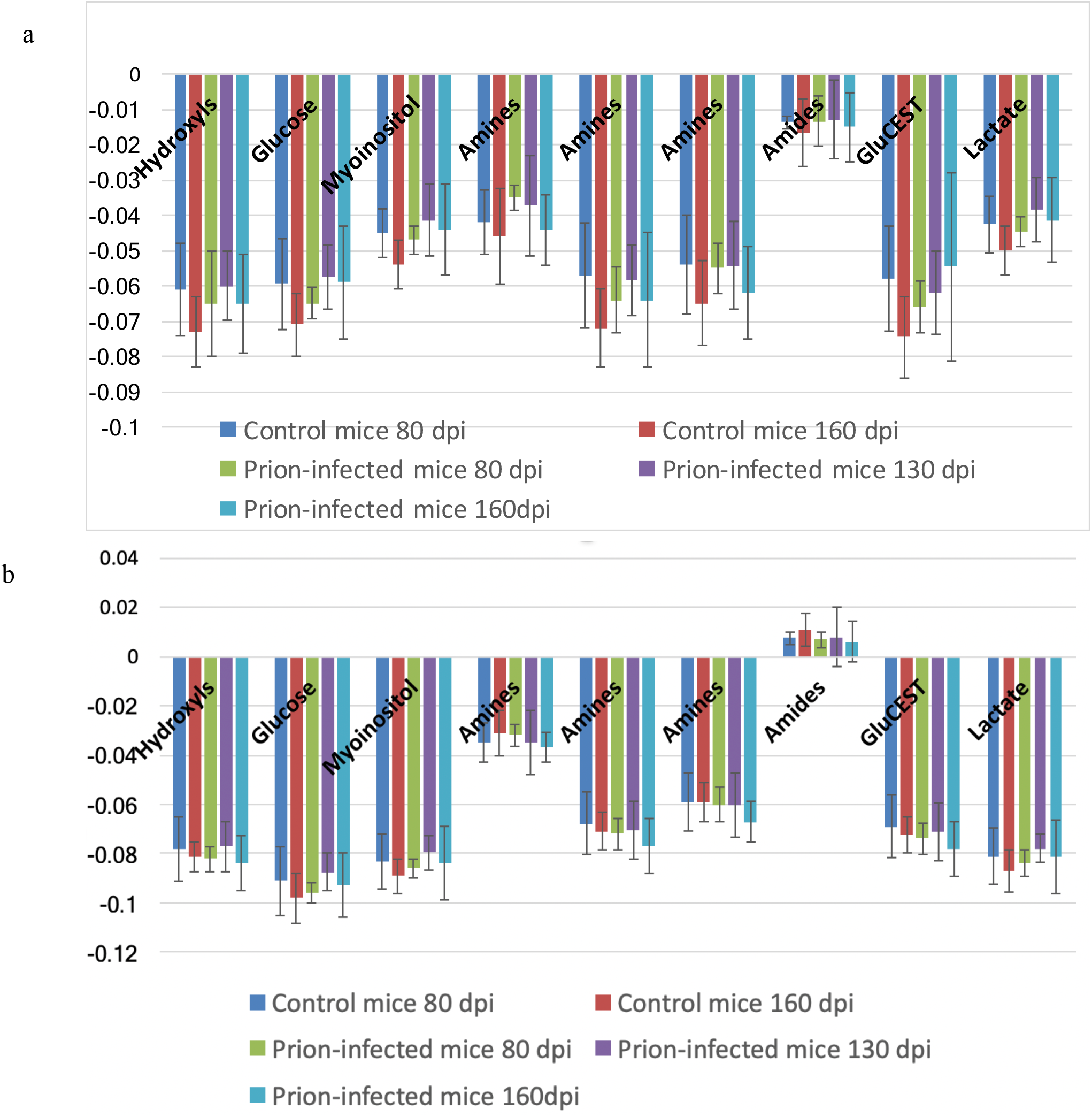
MTR asymmetry values in the cortex of prion and control mice for the frequency offsets reported in Supplementary table 1 at (a) 0.6 μT and (b) 1.2 μT

**Supplementary figure 5:**
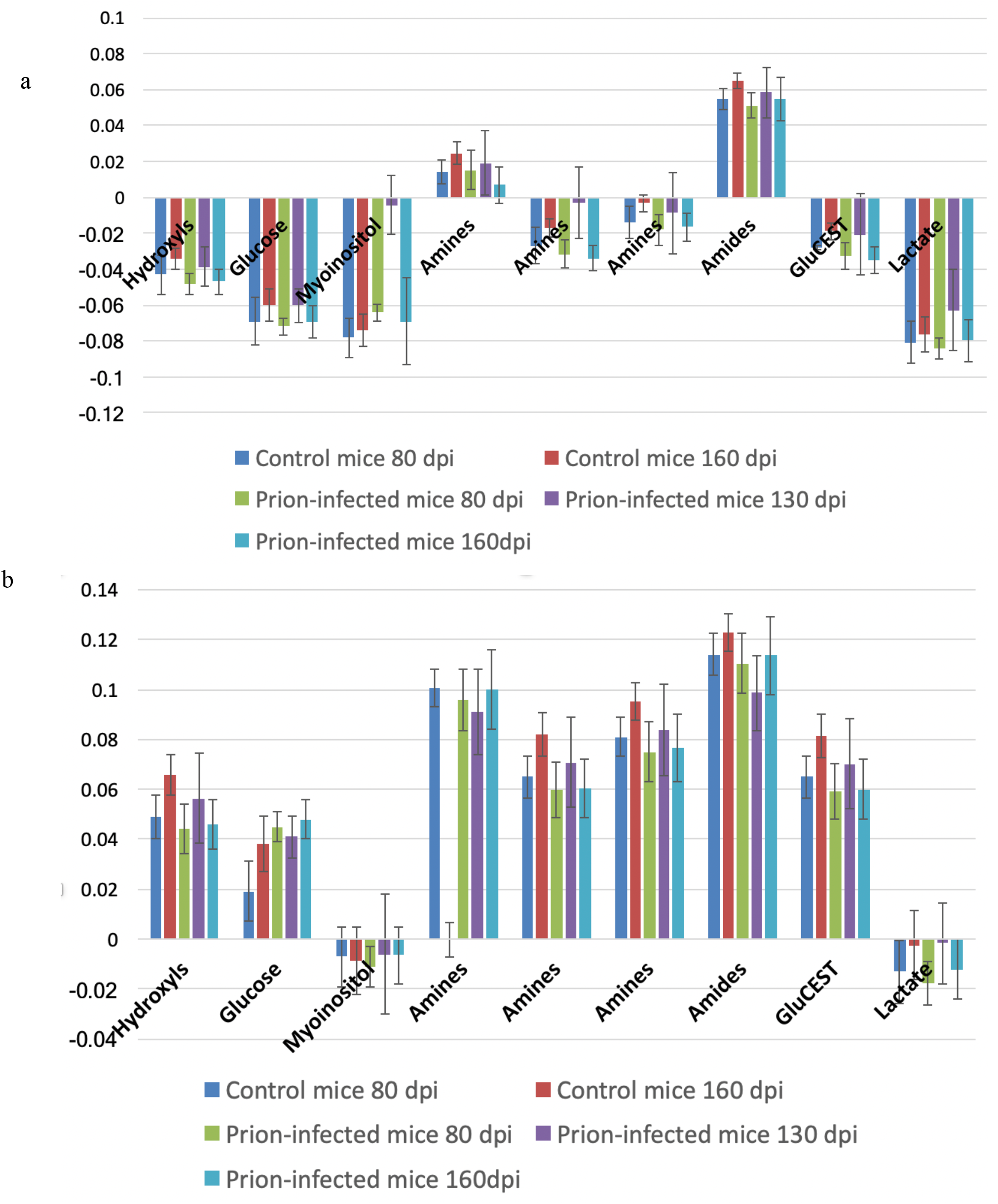
MTR asymmetry values in the cortex of prion and control mice for the frequency offsets reported in Supplementary table 1 at (a) 2.0 μT and (b) 3.6 μT

**Supplementary figure 6:**
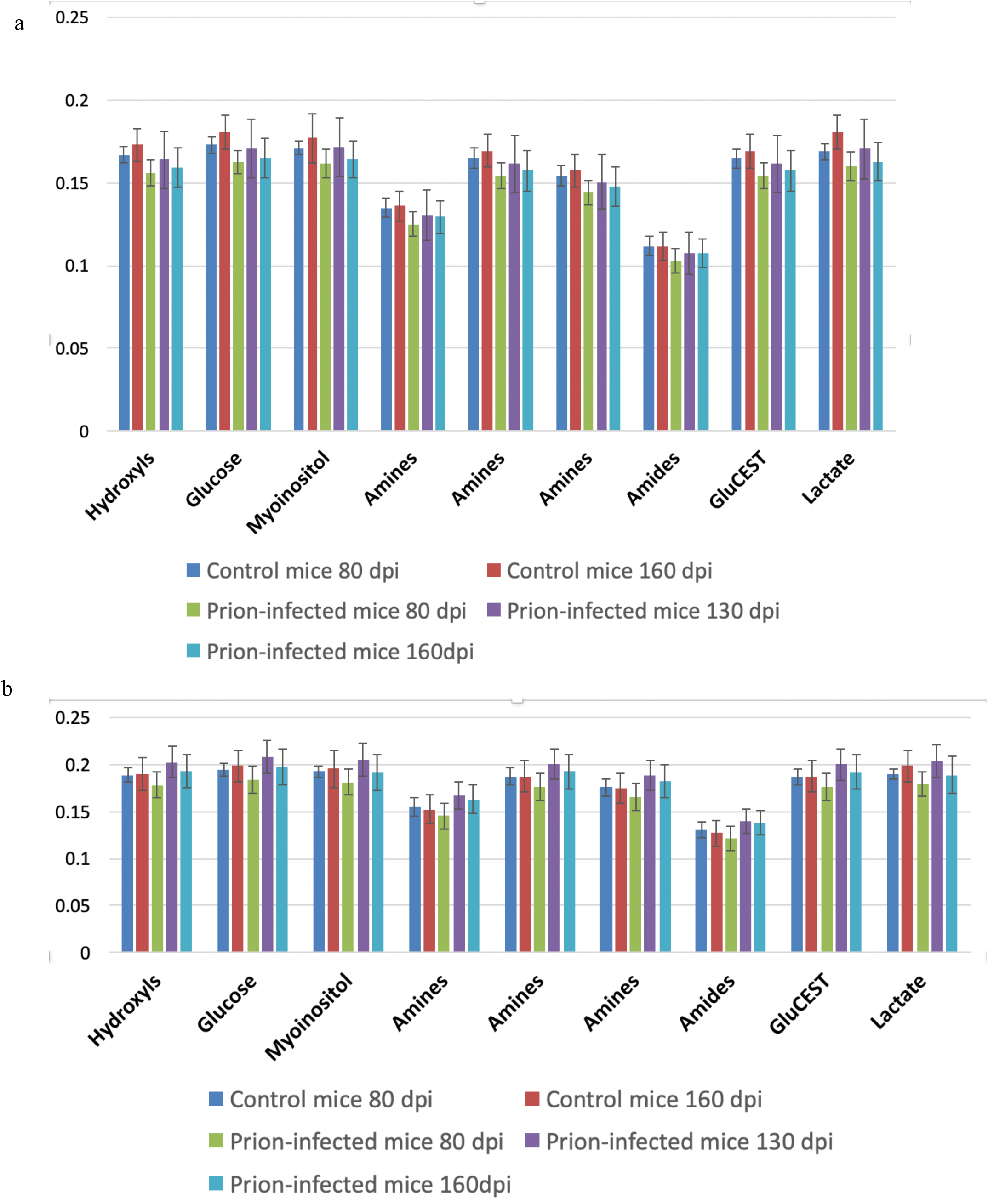
MTR asymmetry values at 10 μT of prion and control mice for the frequency offsets reported in Supplementary table 1 in (a) the cortex and (b) thalamus

**Supplementary figure 7:**
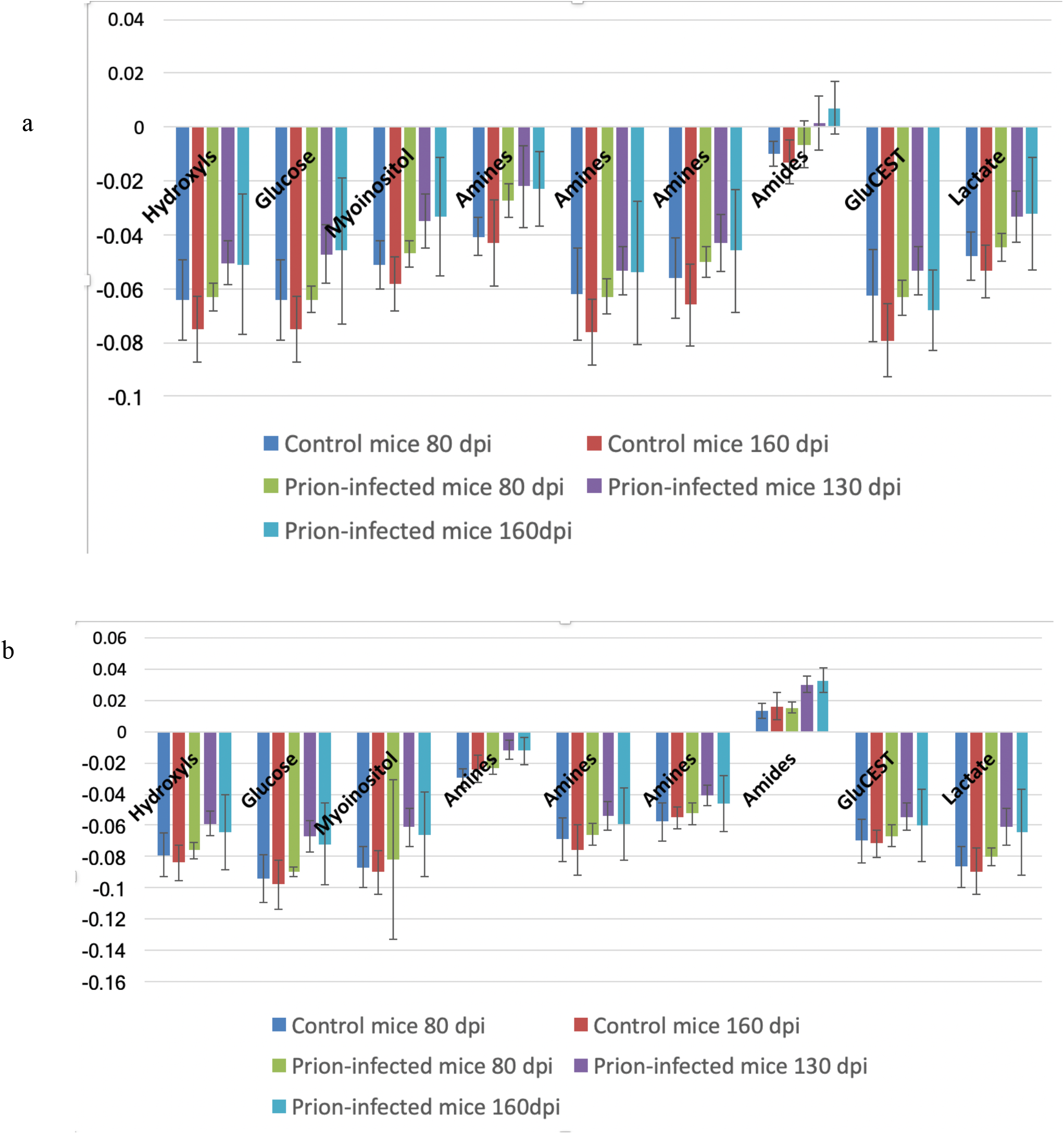
MTR asymmetry values in the thalamus of prion and control mice for the frequency offsets reported in Supplementary table 1 at (a) 0.6 μT and (b) 1.2 μT

**Supplementary figure 8:**
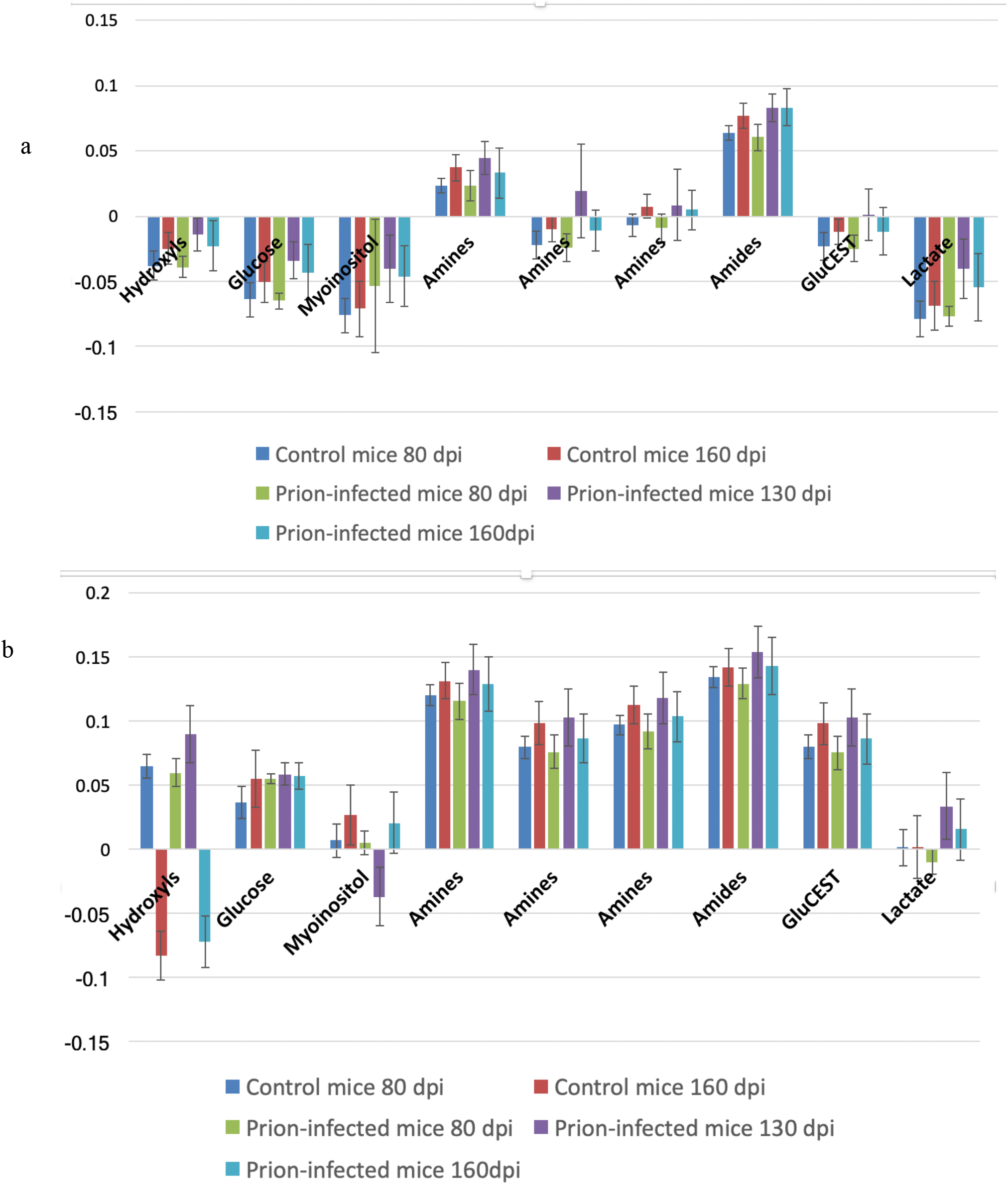
MTR asymmetry values in the thalamus of prion and control mice for the frequency offsets reported in Supplementary table 1 at (a) 2.0 μT and (b) 3.6 μT

**Supplementary Table 2.** displays the significant changes (p values) of MTR asymmetries of 160 dpi prion-infected mice vs age matched controls in thalamus and cortex. Red indicates larger MTR asymmetries in prion-infected mice and grey smaller MTR asymmetries compared to the control group.

Cortex
|  | 0.6 uT | 1.2 uT | 2.0uT | 3.6 uT | 10 uT |
| --- | --- | --- | --- | --- | --- |
| 1 | 0.21 | 0.58 | 0.0015 | 0.0019 | 0.025 |
| 2 | 0.037 | 0.20 | 0.043 | 0.056 | 0.012 |
| 3 | 0.04 | 0.19 | 0.30 | 0.021 | 0.0393 |
| 4 | 0.33 | 0.09 | 0.001 | 0.0046 | 0.12 |
| 5 | 0.32 | 0.04 | 0.019 | 0.005 | 0.06 |
| 6 | 0.17 | 0.13 | 0.00025 | 0.0011 | 0.03 |
| 7 | 0.37 | 0.09 | 0.037 | 0.09 | 0.10 |
| 8 | 0.05 | 0.134 | 0.0002 | 0.001 | 0.003 |
| 9 | 0.06 | 0.19 | 0.29 | 0.03 | 0.004 |

Thalamus
|  | 0.6 uT | 1.2 uT | 2.0 uT | 3.6 uT | 10 uT |
| --- | --- | --- | --- | --- | --- |
| 1 | 0.04 | 0.035 | 0.40 | 0.33 | 0.38 |
| 2 | 0.013 | 0.022 | 0.26 | 0.42 | 0.43 |
| 3 | 0.012 | 0.033 | 0.029 | 0.32 | 0.37 |
| 4 | 0.011 | 0.0127 | 0.27 | 0.40 | 0.10 |
| 5 | 0.038 | 0.12 | 0.35 | 0.17 | 0.20 |
| 6 | 0.04 | 0.07 | 0.44 | 0.11 | 0.303 |
| 7 | 0.00065 | 0.0008 | 0.16 | 0.45 | 0.12 |
| 8 | 0.08 | 0.126 | 0.45 | 0.12 | 0.304 |
| 9 | 0.02 | 0.032 | 0.12 | 0.36 | 0.18 |

**Supplementary Figure 9.**
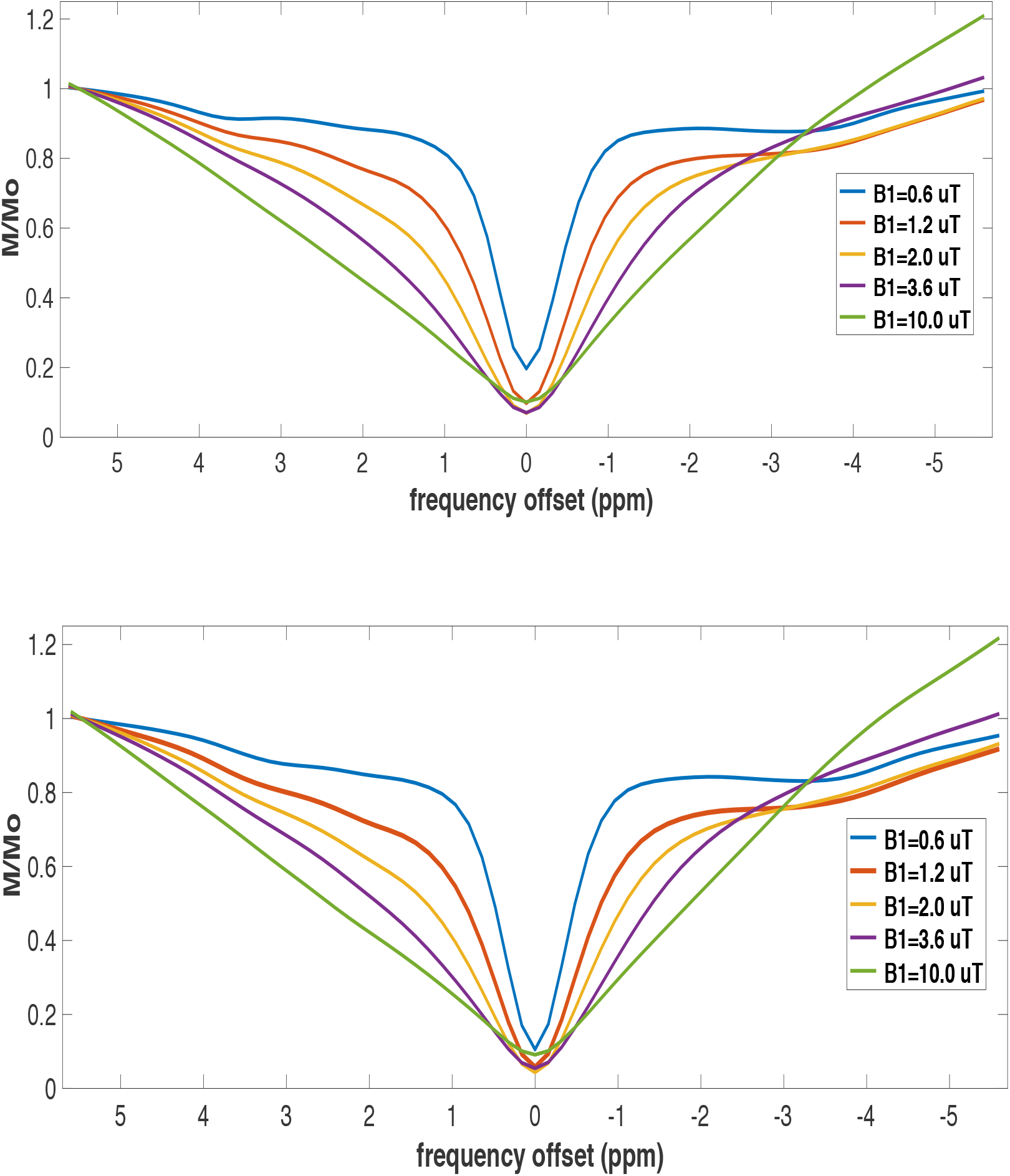
(a) Z-spectra obtained from thalamus of a late stage prion-infected mouse and control mouse (b) at five different irradiation amplitudes.

**Supplementary Figure 10.**
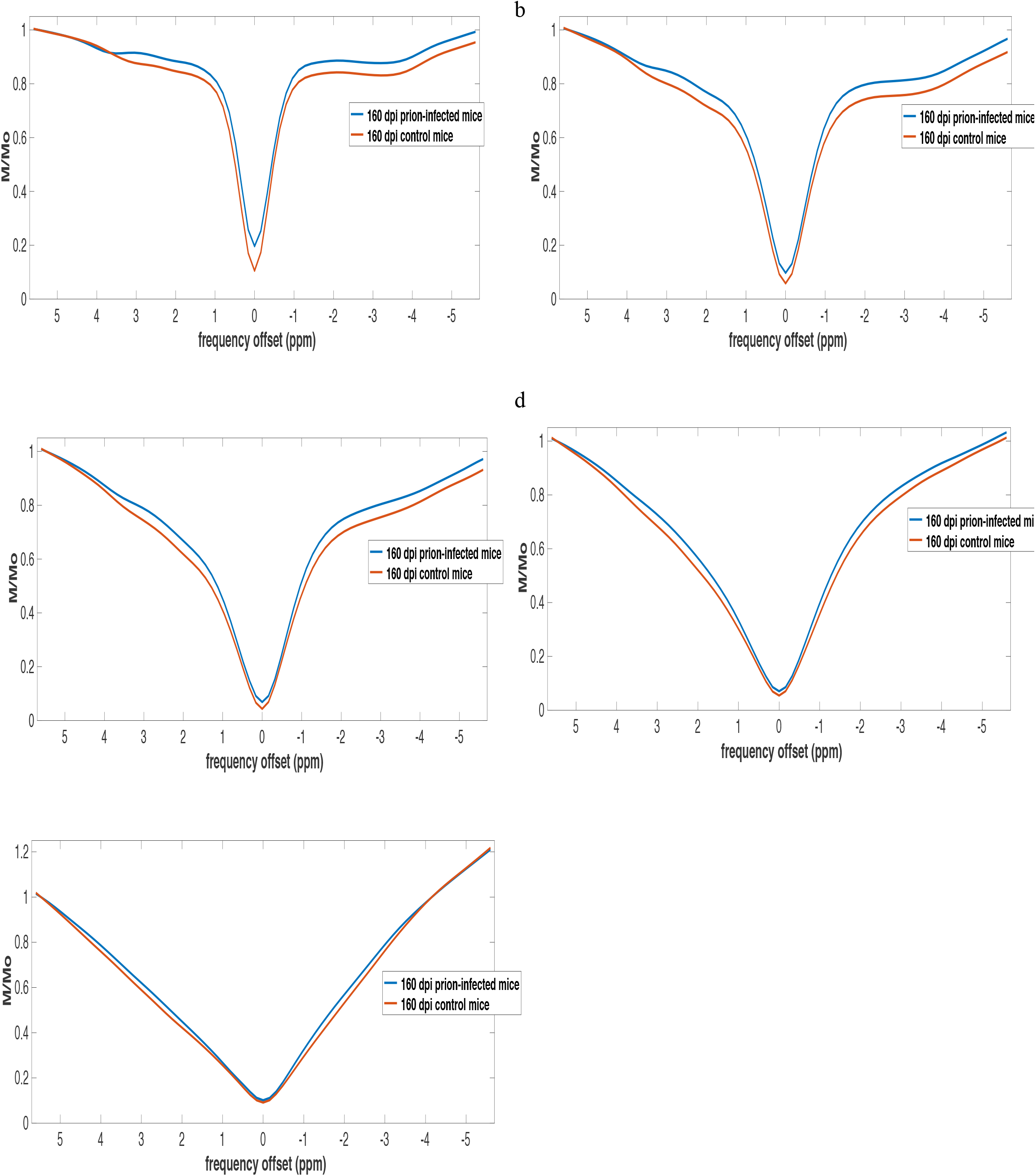
Z-spectra obtained from 160 dpi prion-infected mice and aged matched controls in thalamus at 0.6 **μT** (**a**), 1.2 **μT** (**b**), 2.0 **μT** (**c**), 3.6 **μT** (**d**) and 10 **μT** (**e**).

**Supplementary Figure 11.**
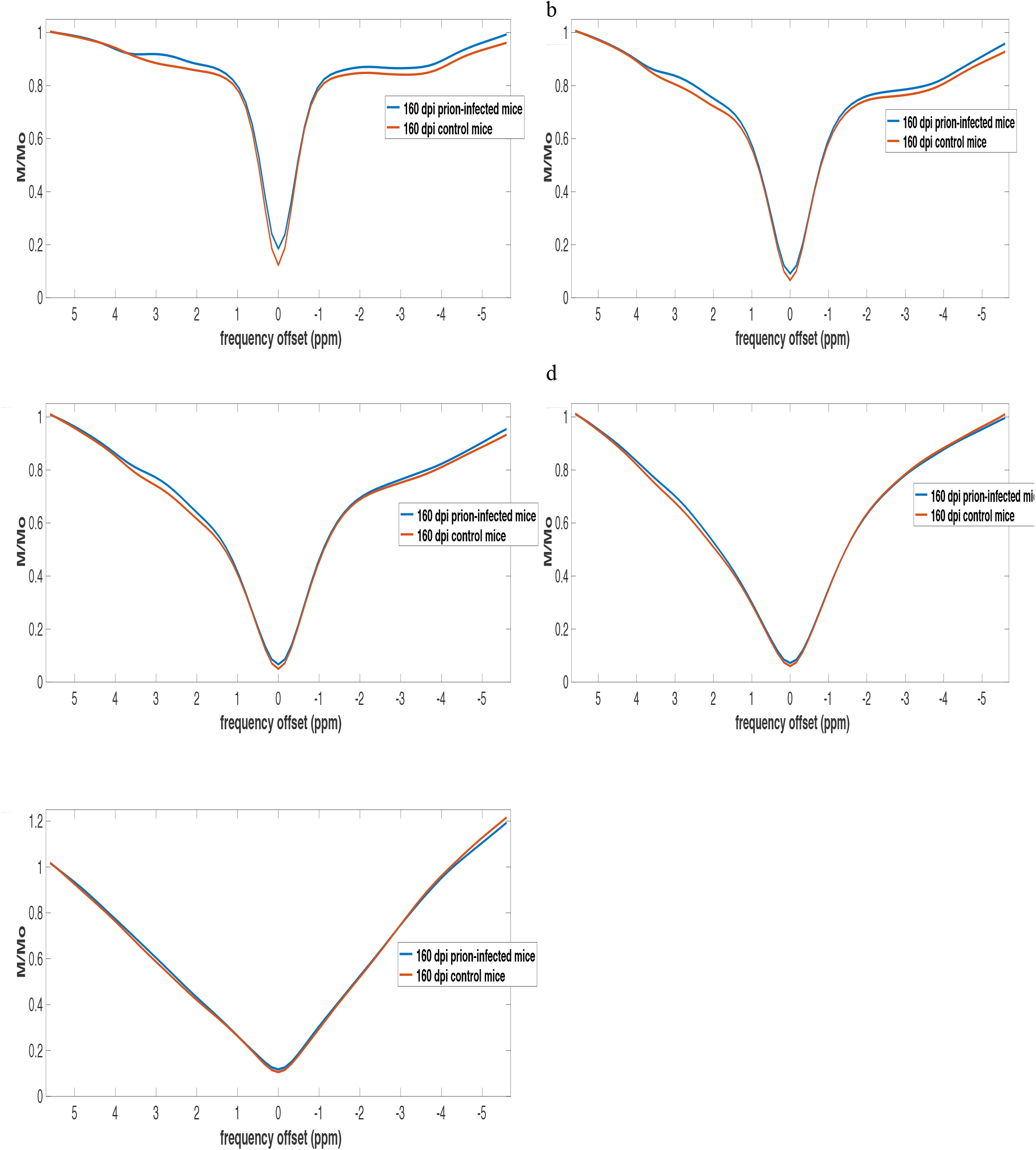
Z-spectra obtained from 160 dpi prion-infected mice and aged matched controls in cortex at 0.6 **μT** (**a**), 1.2 **μT** (**b**), 2.0 **μT** (**c**), 3.6 **μT** (**d**) and 10 **μT** (**e**).

**Supplementary Table 3.** displays the significant changes (p values) of MTR asymmetries of 80 dpi prion-infected mice vs age matched controls in thalamus and cortex. Red indicates larger MTR asymmetries in prion-infected mice and grey indicates smaller asymmetries compared to the control group.

Cortex
|  | 0.6 uT | 1.2 uT | 2.0uT | 3.6 uT | 10 uT |
| --- | --- | --- | --- | --- | --- |
| **1.** | 0.25 | 0.25 | 0.23 | 0.19 | 0.01 |
| **2.** | 0.20 | 0.25 | 0.29 | 0.0029 | 0.01 |
| **3.** | 0.27 | 0.27 | 0.25 | 0.24 | 0.025 |
| **4.** | 0.088 | 0.29 | 0.41 | 0.22 | 0.01 |
| **5.** | 0.43 | 0.41 | 0.23 | 0.16 | 0.01 |
| **6.** | 0.195 | 0.21 | 0.21 | 0.18 | 0.01 |
| **7.** | 0.46 | 0.36 | 0.16 | 0.28 | 0.02 |
| **8.** | 0.17 | 0.22 | 0.20 | 0.18 | 0.02 |
| **9.** | 0.32 | 0.311 | 0.308 | 0.25 | 0.03 |

Thalamus
|  | 0.6 uT | 1.2 uT | 2.0uT | 3.6 uT | 10 uT |
| --- | --- | --- | --- | --- | --- |
| **1.** | 0.46 | 0.30 | 0.42 | 0.25 | 0.07 |
| **2.** | 0.47 | 0.33 | 0.46 | 0.0127 | 0.07 |
| **3.** | 0.23 | 0.20 | 0.35 | 0.17 | 0.06 |
| **4.** | 0.0078 | 0.05 | 0.48 | 0.23 | 0.09 |
| **5.** | 0.20 | 0.22 | 0.39 | 0.22 | 0.08 |
| **6.** | 0.44 | 0.32 | 0.42 | 0.25 | 0.08 |
| **7.** | 0.21 | 0.23 | 0.29 | 0.21 | 0.09 |
| **8.** | 0.46 | 0.32 | 0.42 | 0.25 | 0.08 |
| **9.** | 0.23 | 0.185 | 0.37 | 0.06 | 0.36 |

**Supplementary Figure 12.**
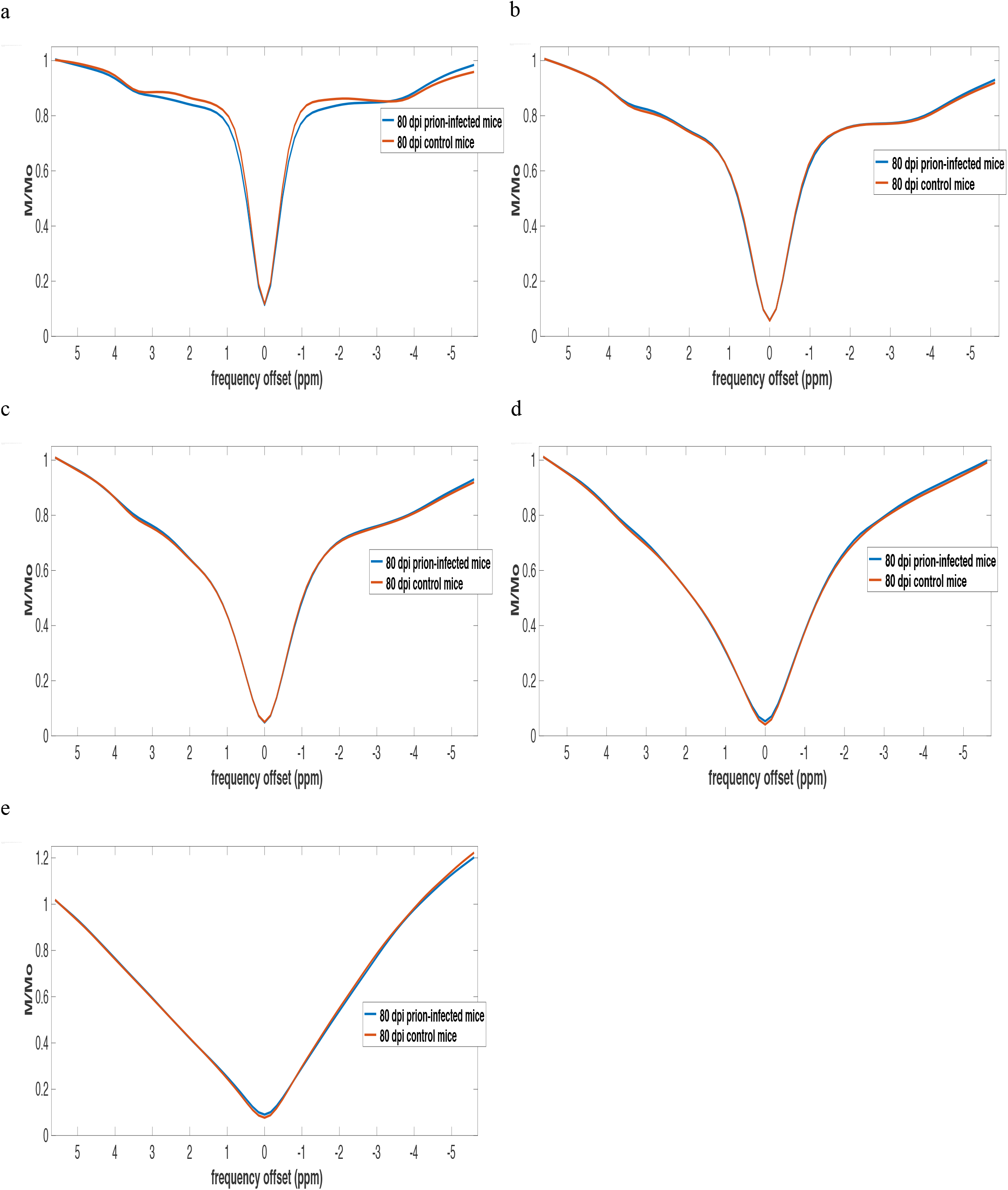
Z-spectra obtained from 80 dpi prion-infected mice and aged matched controls in thalamus at 0.6 **μT** (**a**), 1.2 **μT** (**b**), 2.0 **μT** (**c**), 3.6 **μT** (**d**) and 10 **μT** (**e**).

**Supplementary Figure 13.**
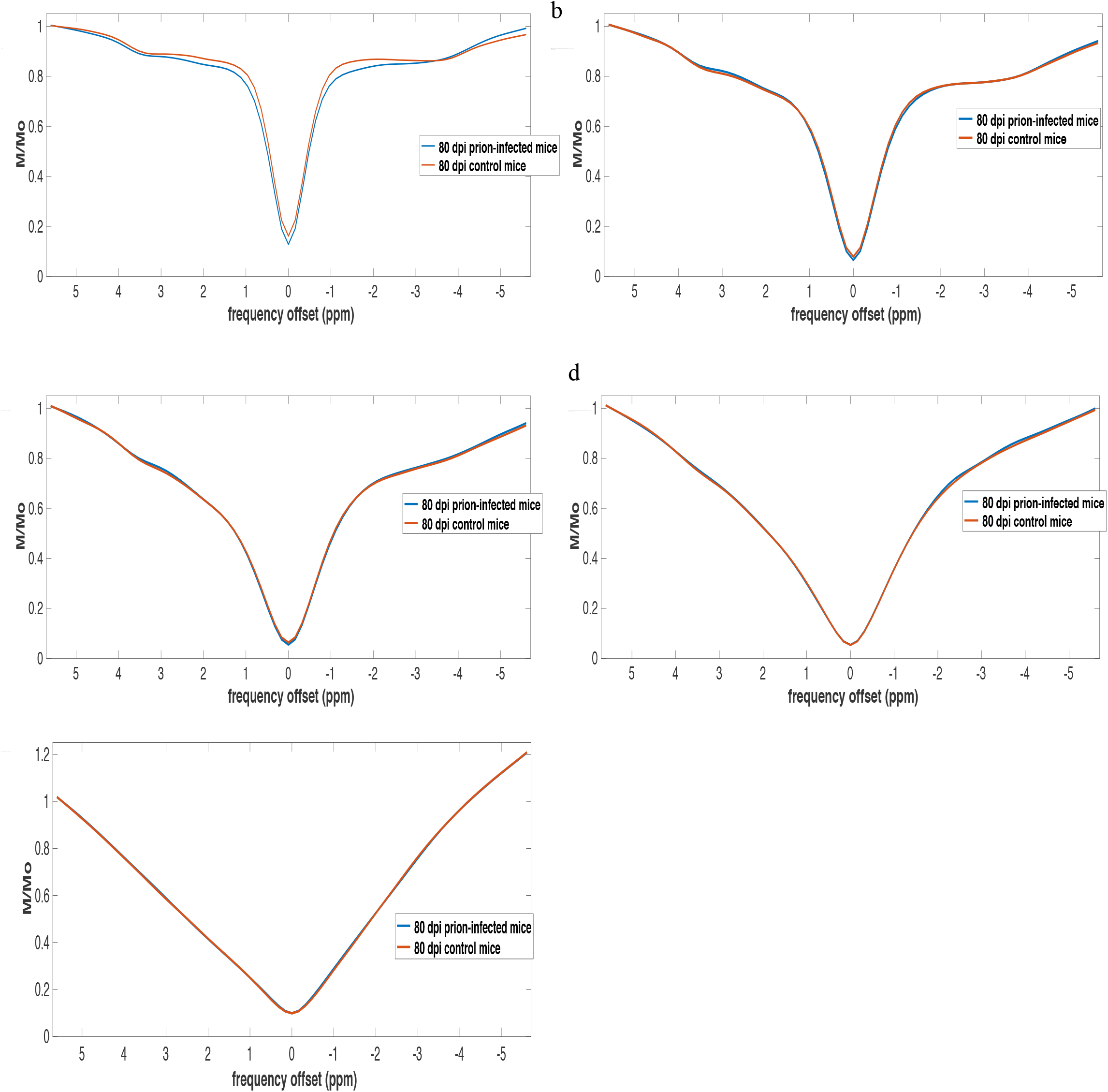
Z-spectra obtained from 80 dpi prion-infected mice and aged matched controls in cortex at 0.6 **μT** (**a**), 1.2 **μT** (**b**), 2.0 **μT** (**c**), 3.6 **μT** (**d**) and 10 **μT** (**e**).

**Supplementary Table 4.**
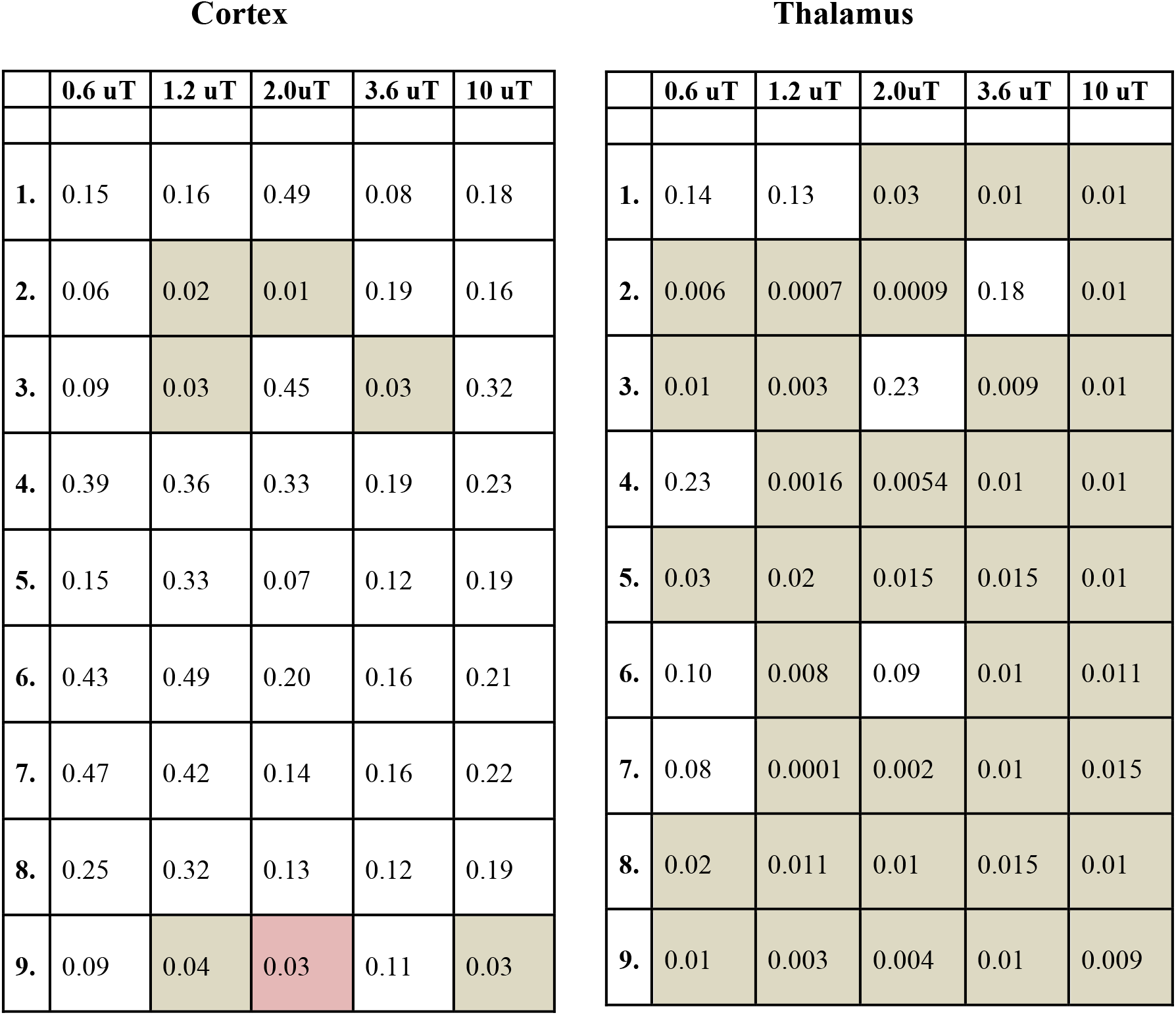
displays the significant changes (p values) of MTR asymmetries of 80 dpi prion-infected mice vs 130 dpi prion-infected mice in thalamus and cortex. Larger MTR asymmetries were detected for the prion-infected group at 130 dpi when compared to 80 dpi prion-infected mice. Comparison was between the prion-infected group 130 dpi vs 80 dpi.

**Supplementary Table 5.** displays the significant changes (p values) of MTR asymmetries of 80 dpi prion-infected mice vs 160 dpi prion-infected mice in thalamus and cortex. Reduced MTR asymmetries were detected for the prion-infected group at 80 dpi when compared to 160 dpi prion-infected mice.

Cortex
|  | 0.6 uT | 1.2 uT | 2.0uT | 3.6 uT | 10 uT |
| --- | --- | --- | --- | --- | --- |
| 1. | 0.47 | 0.37 | 0.49 | 0.33 | 0.32 |
| 2. | 0.18 | 0.28 | 0.22 | 0.27 | 0.33 |
| 3. | 0.25 | 0.32 | 0.40 | 0.17 | 0.32 |
| 4. | 0.03 | 0.06 | 0.11 | 0.31 | 0.22 |
| 5. | 0.49 | 0.19 | 0.25 | 0.40 | 0.31 |
| 6. | 0.13 | 0.08 | 0.41 | 0.38 | 0.27 |
| 7. | 0.38 | 0.39 | 0.27 | 0.34 | 0.15 |
| 8. | 0.16 | 0.20 | 0.28 | 0.45 | 0.31 |
| 9. | 0.27 | 0.34 | 0.22 | 0.32 | 0.17 |

Thalamus
|  | 0.6 uT | 1.2 uT | 2.0uT | 3.6 uT | 10 uT |
| --- | --- | --- | --- | --- | --- |
| 1. | 0.14 | 0.13 | 0.031 | 0.08 | 0.06 |
| 2. | 0.07 | 0.06 | 0.02 | 0.28 | 0.09 |
| 3. | 0.08 | 0.09 | 0.38 | 0.06 | 0.13 |
| 4. | 0.26 | 0.0086 | 0.141 | 0.09 | 0.03 |
| 5. | 0.22 | 0.27 | 0.07 | 0.13 | 0.05 |
| 6. | 0.36 | 0.20 | 0.04 | 0.11 | 0.039 |
| 7. | 0.01 | 0.00032 | 0.0003 | 0.08 | 0.02 |
| 8. | 0.22 | 0.25 | 0.06 | 0.12 | 0.05 |
| 9. | 0.08 | 0.09 | 0.03 | 0.16 | 0.07 |

**Supplementary Figure 14.**
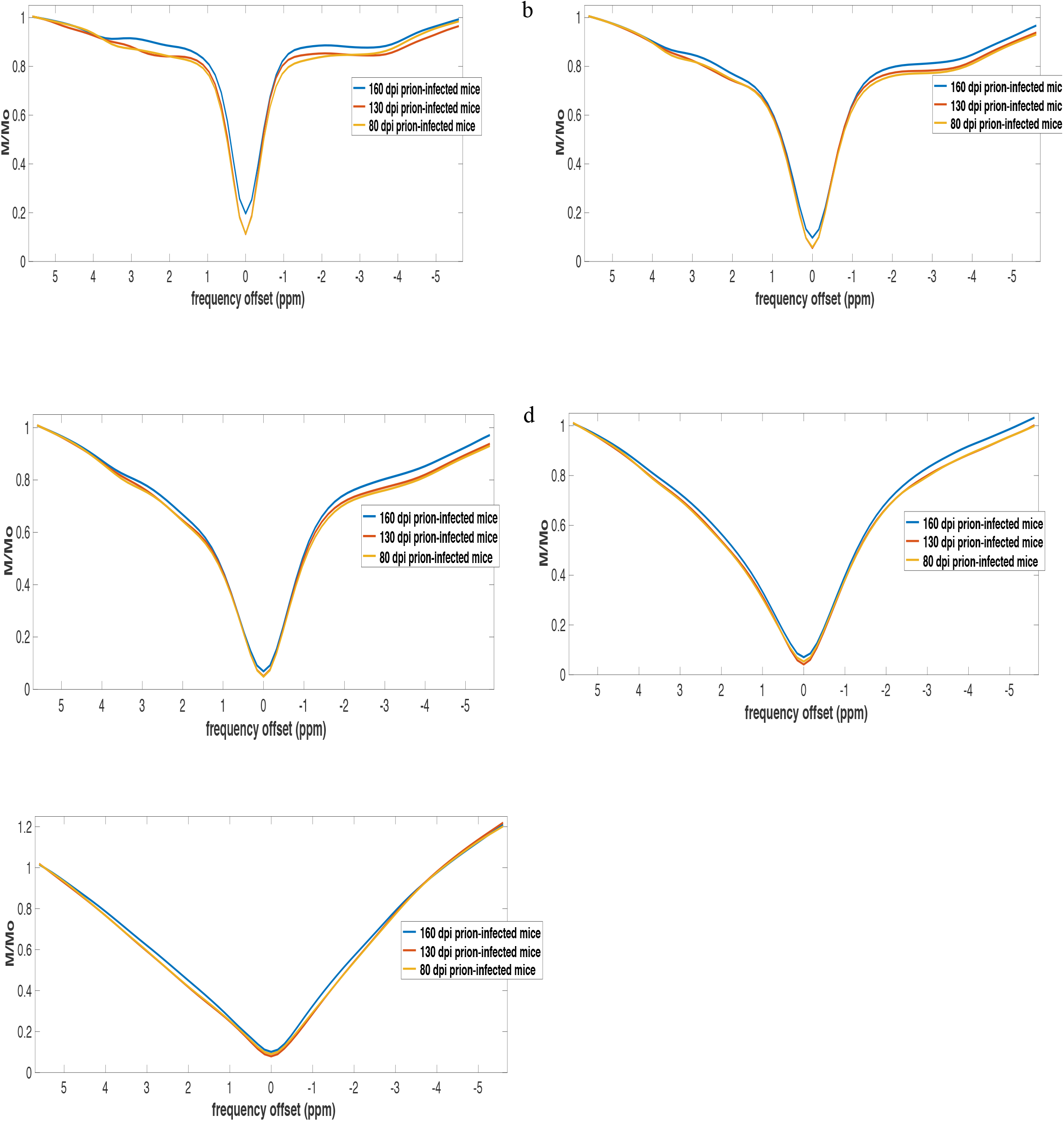
Z-spectra obtained from 160 dpi, 130 dpi and 80 dpi prion-infected mice in thalamus at 0.6 **μT** (**a**), 1.2 **μT** (**b**), 2.0 **μT** (**c**), 3.6 **μT** (**d**) and 10 **μT** (**e**).

**Supplementary Figure 15.**
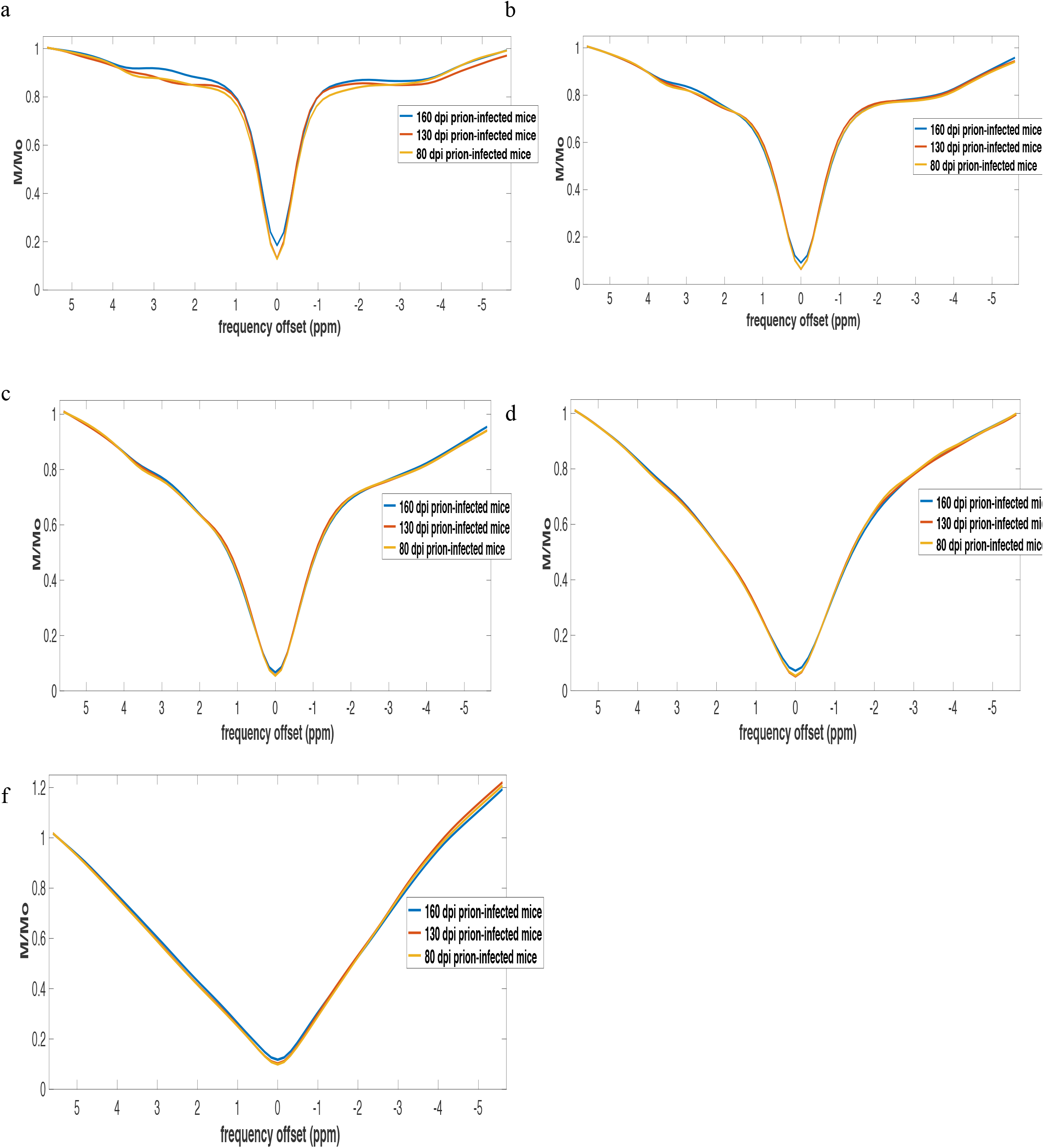
Z-spectra obtained from 160 dpi, 130 dpi and 80 dpi prion-infected mice in cortex at 0.6 **μT** (**a**), 1.2 **μT** (**b**), 2.0 **μT** (**c**), 3.6 **μT** (**d**) and 10 **μT** (**e**).

## Comparison of CEST signal in control mice

### 80 dpi control mice vs 160 dpi control mice

Decreased MTR asymmetries were detected at high irradiation amplitudes (i.e. B_1_ < 2.0 μT) in both thalamus and cortex of 80 dpi mice when compared to 160 dpi control mice. In addition, the MTR asymmetries at 0.6 μT in the cortex of 80 dpi mice were found to be increased compared to 160 dpi control mice. Supplementary Table S5 displays the comparison of MTR asymmetries in 80 dpi and 160 dpi control mice.

**Supplementary Table 6.** displays the significant changes (p values) of MTR asymmetries of 80 dpi vs 160 dpi control mice in thalamus and cortex. Red indicates larger MTR asymmetries in the 80 dpi control mice and grey smaller asymmetries compared to 160 dpi mice.

| Cortex |  |  |  |  |  | Thalamus |  |  |  |  |  |
| --- | --- | --- | --- | --- | --- | --- | --- | --- | --- | --- | --- |
|  | 0.6 uT | 1.2 uT | 2.0uT | 3.6 uT | 10 uT |  | 0.6uT | 1.2uT | 2.0uT | 3.6 uT | 10uT |
| 1. | 0.05 | 0.35 | 0.07 | 0.0077 | 0.12 | 1. | 0.10 | 0.46 | 0.07 | 0.027 | 0.42 |
| 2. | 0.053 | 0.278 | 0.11 | 0.014 | 0.06 | 2. | 0.09 | 0.299 | 0.07 | 0.04 | 0.30 |
| 3. | 0.04 | 0.15 | 0.26 | 0.03 | 0.01 | 3. | 0.15 | 0.36 | 0.14 | 0.06 | 0.165 |
| 4. | 0.38 | 0.26 | 0.011 | 0.011 | 0.428 | 4. | 0.45 | 0.12 | 0.0049 | 0.057 | 0.34 |
| 5. | 0.05 | 0.29 | 0.06 | 0.004 | 0.18 | 5. | 0.07 | 0.39 | 0.03 | 0.0183 | 0.48 |
| 6. | 0.13 | 0.48 | 0.025 | 0.0049 | 0.27 | 6. | 0.23 | 0.36 | 0.0096 | 0.0195 | 0.43 |
| 7. | 0.23 | 0.08 | 0.0085 | 0.05 | 0.46 | 7. | 0.30 | 0.08 | 0.00665 | 0.14 | 0.29 |
| 8. | 0.05 | 0.30 | 0.06 | 0.0049 | 0.17 | 8. | 0.08 | 0.41 | 0.03 | 0.018 | 0.48 |
| 9. | 0.07 | 0.17 | 0.27 | 0.036 | 0.14 | 9. | 0.17 | 0.37 | 0.14 | 0.06 | 0.01 |

## Summary of the mixed effects analysis

**Supplementary table 7.**

|  |  |  |  |
| --- | --- | --- | --- |
| Random-effects GLS regression | Number of obs | = | 66 |
| Group variable: **mouse** | Number of groups | = | 8 |
| R-sq: | Obs per group: |  |  |
| within = **0.5261** | min = |  | 2 |
| between = **0.6345** | avg = |  | 8.2 |
| overall = **0.4944** | max = |  | 10 |
|  | Wald chi2(5) | = | 61768.98 |
| corr(u_i, X) = 0 (assumed) | Prob > chi2 | = | 0.0000 |

| signal | Coef. | Std. Err. | z | P>\|z\| | [95% Conf. Interval] |  |
| --- | --- | --- | --- | --- | --- | --- |
| power | .0387265 | .0003435 | 112.74 | 0.000 | .0380532 | .0393998 |
| age |  |  |  |  |  |  |
| 130 | .0075413 | .0047406 | 1.59 | 0.112 | -.00175 | .0168326 |
| 160 | .0151419 | .0034283 | 4.42 | 0.000 | .0084226 | .0218612 |
| disease_cat |  |  |  |  |  |  |
| prion | -.0184813 | .0034094 | -5.42 | 0.000 | -.0251636 | -.011799 |
| rio2_cat |  |  |  |  |  |  |
| thalamus | -.0046194 | .0030801 | -1.50 | 0.134 | -.0106563 | .0014175 |
| _cons | 0 | (omitted) |  |  |  |  |

